# Identifying the Copper Coordination Environment Between Interacting Neurodegenerative Proteins: A New Approach Using Pulsed EPR with ^14^N/^15^N Isotopic Labelling

**DOI:** 10.1101/2024.11.14.623671

**Authors:** Amanda Smart, Kevin Singewald, Zikri Hasanbasri, R. David Britt, Glenn L. Millhauser

## Abstract

The trafficking and aggregation of neurodegenerative proteins often involves the interaction between intrinsically disordered domains, stabilized by the inclusion of physiologic metal ions such as copper or zinc. Characterizing the metal ion coordination environment is critical for assessing the stability and organization of these relevant protein-protein interactions but is challenging given the lack of regular molecular order or global structure. The cellular prion protein (PrP^C^) binds both monomers and aggregates of the Alzheimer’s amyloid-beta peptide (Aβ), promoting interactions of relevance to Aβ internalization across the cellular plasma membrane and aberrant signaling in neurodegenerative disease, respectively. Both proteins bind Cu^2+^ with high affinity, suggesting the existence of a ternary complex with copper bridging between the two proteins through His coordination. In this work, we describe a novel approach utilizing multiple EPR experiments to characterize the simultaneous Cu^2+^ coordination of PrP^C^ and Aβ. Uniformly ^15^N-labeled PrP^C^ is used in conjunction with natural abundance ^14^N Aβ, the combination of which leads to distinct energy manifolds for paramagnetic Cu^2+^ and resolved by the pulsed EPR experiments ESEEM and HYSCORE. We develop acquisition parameters to simultaneously optimize ^14^N (I = 1) and ^15^N (I = ½) pulsed EPR signals and we also advance the theory of ESEEM and HYSCORE to quantitatively describe multiple ^15^N imidazole coordination. Together, these findings provide a detailed view of how Cu^2+^ bridges between the two proteins in this complex, along with a global strategy for assessing the copper environment with other interacting neurodegenerative proteins.

## Introduction

Neurodegenerative diseases are initiated by protein-protein interactions, typically in the form of extracellular aggregates. In turn, these aggregates trigger downstream events ultimately compromising neuronal function. Alzheimer’s disease (AD) begins with intermolecular association of Aβ, a 40-42 amino acid peptide derived from proteolytic cleavage of the amyloid precursor protein (APP) (1). Interestingly, the cellular prion protein (PrP^C^), which itself causes a separate set of neurodegenerative diseases, is recognized as a high affinity receptor for both monomeric Aβ and oligomeric (Aβo) (2–6). Monomeric Aβ binding to PrP^C^ results in the translocation of Aβ through endocytosis, possibly leading to its intracellular accumulation (4, 7). Aβo binding to PrP^C^ is proposed to trigger aberrant transmembrane signaling through the metabotropic glutamate receptor 5 (mGluR5) ultimately causing intracellular tau aggregation and the formation of neurofibrillary tangles (6, 8, 9). Metal ion binding is likely to play a significant role in all these processes. Senile plaques formed primarily by Aβ aggregates are rich with both copper and zinc (10, 11). The occlusion of these metal ions is proposed to both stabilize aggregate structure and contribute to deleterious chemical reactions through the formation of reactive oxygen species (12–16). In their monomeric forms, both Aβ and PrP^C^ bind Cu^2+^ with high affinity (10, 15, 17). As such, assessment of the interaction between these two proteins must consider their respective metalloprotein characteristics.

Murine PrP^C^ is composed of 208 amino acids (residues 23–230), post-translationally modified with a C-terminal glycophosphatidylinositol anchor and two Asn-linked glycans at residues 180 and 196. The N-terminal segment (residues 23–125, after removal of the signal peptide) is an intrinsically disordered peptide (IDP), whereas the C-terminal domain is primarily helical. Within the N-terminal IDP is the octapeptide repeat (OR) domain, residues 59 to 90, composed of the sequence (PHGGXWGQ)_4_ in mouse PrP^C^ (where X is Gly in repeats one and four, Ser in repeats two and three), which binds up to four equivalents of Cu^2+^ (18, 19). It is now established that Cu^2+^ forms a coordination bridge between the OR domain and the C-terminal domain; the stability of this bridge is further enhanced by glycosylation (20–22). Without this critical N-terminal—C-terminal interaction, the N-terminal domain is untethered and thus drives highly cytotoxic membrane leakage (23).

We demonstrated in previous work that PrP^C^ translocates monomeric Aβ across the plasma membrane by endocytosis (4). Assays were performed with both Aβ(1–40) and Aβ(1-30), the latter of which lacks the peptide’s hydrophobic C-terminal segment and thus remains soluble. By switching Aβ stereochemistry, we showed that its translocation requires a direct protein-protein interaction. Aβ(1-30) is flexible, possesses three His residues within its first 14 amino acids and, similar to PrP^C^, avidly binds Cu^2+^ (4, 16).

Given the demonstrated interaction between PrP^C^ and Aβ and the fact that both are rich with His residues, we propose that Cu^2+^ promotes intermolecular coordination linkages. However, assessing metal ion coordination involving IDP segments in protein complexes is refractory to traditional methods that depend on well-defined protein structure or long-range order. Traditionally, the EPR methods of electron spin echo envelope modulation (ESEEM) (24) and hyperfine sublevel correlation (HYSCORE) (25) are ideal for assessing His coordination (26–28), however, they cannot distinguish which protein is presenting a specific His residue. Here we develop a new strategy for determining how Cu^2+^ stabilizes intermolecular interactions using Aβ-PrP^C^ as a model system. Our approach combines the pulsed EPR techniques of ESEEM and HYSCORE along with ^15^N isotopic labeling of PrP^C^, thus enabling the assignment of spectroscopic signals to residues of the specific interacting partner. We develop data acquisition parameters to produce high quality spectra simultaneously from ^14^N and ^15^N pulsed EPR signals. We also expand the theory of ^15^N pulsed EPR to account for multiple His residue coordination. Our results not only provide a compelling model for how Cu^2+^ stabilizes the interaction between Aβ and PrP^C^, but they also yield a systematic EPR methodology for assessing Cu^2+^ coordination in more general protein complexes.

## Experimental procedures

### Protein Expression and Purification

The prion protein was expressed as previously described (29). In brief, the *M. musculus* PrP(23–230) construct cloned into the pJ414 vector (DNA 2.0) was transformed and expressed using *E. coli* (BL21 (DE3) Invitrogen). The protein was then purified following established methods (30). Bacteria were grown in M9 minimal media supplemented with ^15^N ammonium chloride (1 g/L) (Cambridge Isotopes) for uniformly ^15^N-labeled protein. Cells were grown at 37 °C until an OD_600_ of 1.0 was reached. Then, 1 mM isopropyl β-D-1-thiogalactopyranoside (IPTG) was added and cells were grown at 25 °C overnight. Proteins were extracted from inclusion bodies at room temperature with 8 M guanidinium chloride (GndHCl), 100 mM Tris, and 100 mM sodium acetate at pH 8. The soluble fraction was purified by Ni^2+^-immobilized metal-ion chromatography (IMAC). The column was washed and proteins were eluted with 5 M GndHCl, 100 mM Tris, and 100 mM sodium acetate at pH 4.5. The partially purified proteins were brought to pH 8 with 6 M potassium hydroxide (KOH) and left at 4 °C for 2 days to oxidize the native disulfide bridge. Finally, the proteins were desalted into 50 mM potassium acetate buffer and purified on C8 column with reverse-phase liquid chromatography. The purified PrP^C^ was lyophilized and stored at -70 °C.

Aβ30 was transformed into a pET-28b(+) (Novagen) vector expressing His-tagged small ubiquitin-like modifier (His-SUMO) (Obtained from Carrie Partch at UCSC) with Gibson cloning as previously described (31). Primers were purchased form Invitrogen to linearize His-SUMO DNA and create a linear fragment of Aβ30 DNA in pJ414 using Phusion High-Fidelity PCR Master Mix (New England Biolabs). Linearization reactions were run on a 1% agarose gel and extracted with GeneJET Extraction Kit (Thermo Fisher Scientific). Gibson reactions were run using Gibson Assembly Master Mix (New England Biolabs) and transformed into *E. coli* (DH5ɑ (DE3) Invitrogen). Colonies were grown and pure DNA was extracted using the Qiagen Mini prep kits and verified by DNA sequencing.

The construct was then transformed and expressed in *E. coli* (BL21 (DE3); Invitrogen). Cells were grown in Luria broth media (Research Product International) at 37 °C to an OD_600_ of 0.6-0.8. Then, 1 mM IPTG was added, and the cells continued to grow overnight at 18 °C. Next, the cells were harvested using a Sorvall Lynx 6000 centrifuge at 4 °C and 4000 rpm. Cells were resuspended in lysis buffer (50 mM Tris, 300 mM sodium chloride (NaCl), 1 mM β-mercaptoethanol (βME), and Pierce Protease Inhibitor Tablets (Thermo Fisher Scientific), pH 7.5). Afterwards, cells were sonicated using a FB505 Ultrasonic Processor with 15 second on, 30 second off pulses for 5 minutes. Cells were then centrifuged at 17,000 rpm for 45 minutes and purified using IMAC at 4 °C. The column was washed with 60 mL of Lysis buffer, followed by 30 mL of cleavage buffer (50 mM Tris, 50 mM NaCl, 30 mM Imidazole, 1 mM, pH 7.5). Proteins were incubated on column with 3 mM recombinantly expressed His-Ubl-specific protease 1 overnight at 4 °C. Protein was then eluted from the column with elution buffer (50 mM Tris, 50 mM Imidazole, βME, pH 7.5) and purified using HPLC on an Agilent PLRP-S column under basic conditions as previously described (4). Protein mass was confirmed using a SCIEX ExionLC liquid chromatography system in conjunction with a SCIEX X500B QTOF mass spectrometer, Figure S1. Lastly, the purified Aβ30 was lyophilized and stored at -70 °C.

### Electron Paramagnetic Resonance

Lyophilized ^15^N-PrP^C^ was suspended in water, where Aβ30 was suspended in a small amount of 20 mM KOH and then diluted with water. The concentrations were calculated from the absorbance at 280 nm with an ε of 63,495.0 M^-1^•cm^-1^ and 1490.0 M^-1^•cm^-1^ for ^15^N-PrP^C^ and Aβ30 respectively. All samples were made to 100 μM for each protein, 100 µM Cu^2+^, 25 mM 4-(2-hydroxyethyl)-1-piperazineethanesulfonic acid (HEPES) buffer with 20% glycerol as a cryoprotectant (32), and the pH was adjusted to 7.4 with 100 mM KOH. After samples sat at room temperature for at least 15 minutes they were frozen in 4 mm quartz X-band tubes (Wilmad-LabGlass). For samples at pH 3.5, the 25 mM HEPES was substituted for 25 mM potassium acetate and the pH was adjusted with 100 mM HCl.

Continuous wave (CW)-EPR spectra were recorded at 121 K with a Bruker E500 CW-EPR spectrometer operating at X-band frequency (∼9.3 GHz) using an ER4122SHQE resonator (Bruker). CW-EPR data was simulated using EasySpin 5.1.10 toolbox within the MATLAB (The Mathworks Inc., Natick, MA) R2022b software suite (33).

Electron Spin Echo Envelope Modulations (ESEEM) and Hyperfine Sublevel Correlation Spectroscopy (HYSCORE) time domain signals were acquired on an X-band (∼9.7 GHz) Bruker E580 EPR spectrometer using a MD4 resonator. Pulsed experiments were performed at the maximum of the Cu^2+^ spectrum, 3316 G. Furthermore, the experimental temperature was set to 18 K to optimize the signal to noise of Cu^2+^ (34). The ESEEM pulse sequence was π/2-τ-π/2-T-π/2-τ-echo (24) with four-step phase cycling. Here, we set τ to the hydrogen blind spot, which was calculated to be 210 ns for the magnetic field used (24, 35). The initial T values was set to 12 ns and incremented by 16 ns for 602 points and the π/2-pulse was 8 ns. Experiments using other T and τ values can be found in Figure S7. After acquisition, the ESEEM time domain was intensity normalized, fit to an exponential decay for background subtraction, zero-filled to 2048 points, and Fourier transformed using Bruker XEPR software. HYSCORE spectra were obtained using the pulse sequence π/2-τ-π/2-t_1_-π-t_2_-π/2-τ-echo (25). Here, τ was set to 210 ns to be consistent with the ESEEM experiments, π-pulse was 16 ns, and t_1_ and t_2_ were initially 40 ns with 32 ns increments for 256 points each. The HYSCORE spectra were collected using a four-step phase cycling. The resulting HYSCORE time domain signal was analyzed using Hyscorean (36). The time domain signals utilized Chebyshev apodization and were zero-filled to 1024×1024 points. Lastly, the HYSCORE spectra were obtained with a 2D Fourier transformed. Simulation of the HYSCORE spectra were obtained using the EasySpin toolbox in MATLAB R2022b (33, 37).

## Results and Discussion

### Copper coordination to PrP^C^ and Aβ

The linear sequences of Aβ and PrP^C^ are shown in Figure 1A. Aβ is a 40-42 amino acid long IDP that quickly aggregates, making it difficult to study in biophysical assays. To overcome this tendency to aggregate, a truncated form of Aβ, consisting of the first 30 residues, is used. This truncated Aβ, referred to as Aβ30, is the longest non-aggregating fragment found that retains the ability to bind to both Cu^2+^ and PrP^C^ (4). PrP^C^ has an intrinsically disordered N-terminal domain (residues 23-120) and a structured C-terminal domain (residues 121-230). The flexibility of the N-terminal domain allows for the protein to interact with multiple binding partners, as well as fold back onto the structured domain in the presence of Cu^2+^(29, 30, 38–41). Within the N-terminal domain of PrP^C^, there are two proposed Aβ binding domains at residues 23-31 and 95-115 (2, 8, 42, 43) as well as the OR domain, a 4x repeated sequence each containing a metal binding histidine (17, 18). The OR domain can bind up to four Cu^2+^ ions, however PrP^C^ can bind up to six equivalents of Cu^2+^ simultaneously with additional N-terminal sites at H95 and H110 (44, 45). In Figure 1B we show the coordination of Aβ to Cu^2+^ at physiological pH. This coordination environment consists of the N-terminal amine, a backbone carbonyl, and histidine at residue 6 and residue 13 or 14 (10, 46). Although PrP^C^ can bind up to six copper ions at once, conditions used in this work, i.e., one molar equivalence of Cu^2+^ to PrP^C^ at physiological pH results in one Cu^2+^ per PrP^C^ (18). The coordination of PrP^C^ to Cu^2+^ under these conditions consists of four histidine residues, one from each His in the OR domain, shown in Figure 1C. It should be noted that the binding affinity of Cu^2+^ for PrP^C^ is about 30-fold greater than that of Aβ30. Specifically, as calculated from the glycine competition assay in Figure S2, the binding affinities of Cu^2+^ to Aβ30 and PrP^C^ under the buffer conditions in this study are 16.0 ± 14 nM and 0.519 ± 0.32 nM, respectively. These values fall well within the range of previous work (16, 17).

**Figure 1.**
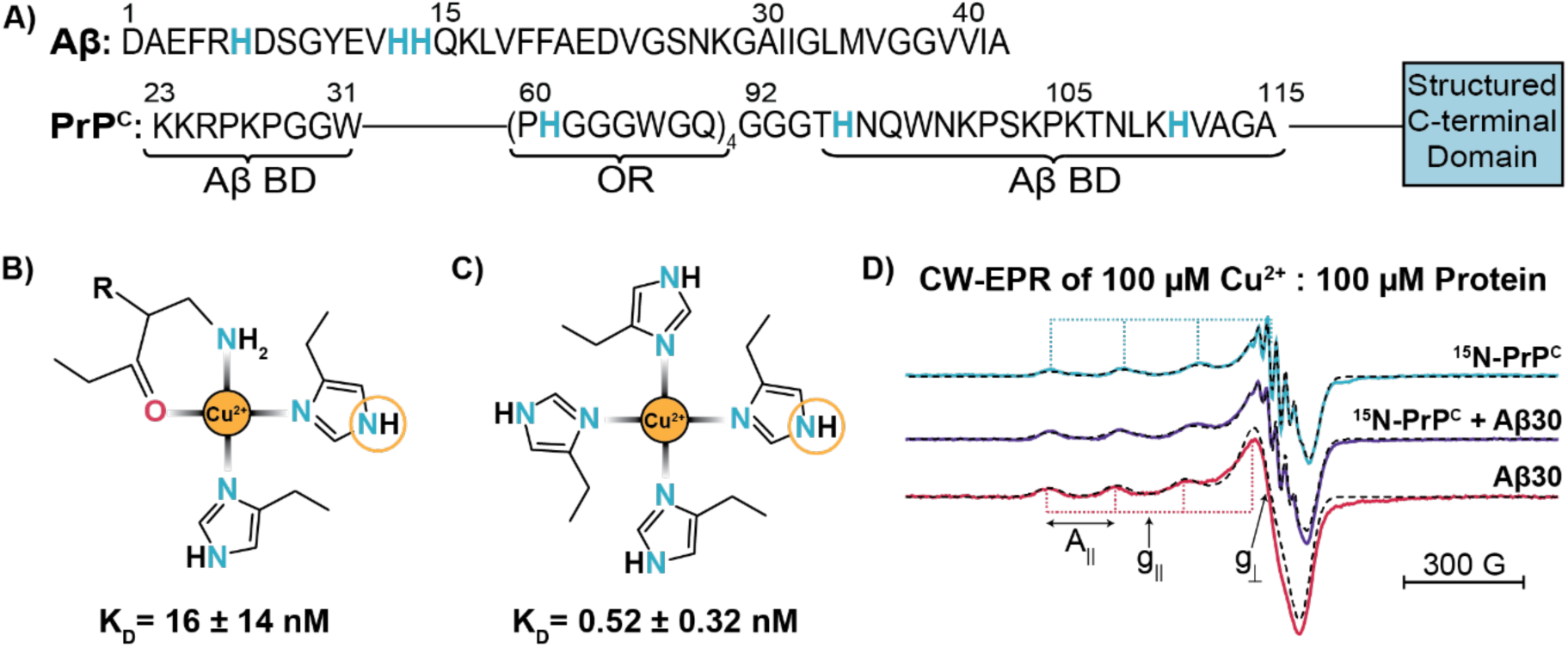
Continuous-wave EPR spectra of Cu^2+^ coordinating to Aβ30 and PrP^C^. A) The linear sequence for Aβ (top) and PrP^C^ (bottom) are shown. Histidine residues that participate in Cu^2+^ binding are blue and bolded. Both Aβ and the N-terminus of PrP^C^ (residues 23-125) are intrinsically disordered. The C-terminus of PrP^C^ (residues 125-230) is structured and represented as the blue box. The two putative Aβ binding domains (Aβ BD) and Cu^2+^ binding domain known as the octapeptide repeat (OR) domain are labeled in PrP^C^. B) The consensus Aβ-Cu^2+^ coordination environment at physiological pH consists of two histidine residues, a backbone carbonyl, and the N-terminal amine. C) The PrP^C^-Cu^2+^ coordination at physiological pH and at equimolar ratio consist of four histidine residues, all from the OR domain. The corresponding affinity (reported as K_D_ values) for the shown coordination state is listed below B) and C). The circled nitrogen on the histidine, known as the remote nitrogen or distally coupled nitrogen, is monitored by ESEEM and HYSCORE EPR. D) The stacked CW of ^15^N-PrP^C^ (blue), ^15^N-PrP^C^:Aβ30 (purple) and Aβ30 (red) are all measured with one equivalent copper at pH 7.4. The sharp, high field features are due to superhyperfine splitting from directly coordinated ^15^N. The dashed spectra overlayed are the respective simulated spectrum. A_∥_ is shown as the separation between the low field peaks, indicated by the blue (PrP^C^) and red (Aβ30) dotted lines. The field related to g_∥_ and g_⊥_ are labeled.

For the following continuous wave (CW) X-band EPR experiments, PrP^C^ was isotopically labeled with ^15^N, whereas Aβ30 was expressed without isotopic labeling. The use of ^15^N or natural abundance does not influence the copper A_∥_ and g_∥_ values obtained from CW-EPR, as shown in Figure S3. In Figure 1D, the top trace is ^15^N-PrP^C^ with A_∥_ = 174 G and g_∥_ = 2.252. The bottom trace is Aβ30 with A_∥_ = 163 G and g_∥_ = 2.269. Both spectra report similar A_∥_ and g_∥_ values to previous work (10, 18, 47). Notably, the ^15^N-PrP^C^ spectrum represents a highly homogeneous Cu^2+^ coordination environment that reveals superhyperfine splitting in the perpendicular spectral region. By comparison, the Aβ30 spectra shows no superhyperfine splitting. Finally, the middle trace is the combination sample containing ^15^N-PrP^C^, Aβ30 and Cu^2+^ at equimolar ratios where A_∥_ = 171 and g_∥_ = 2.263. Consideration of the affinities from the glycine competition assay, Cu^2+^should preferentially coordinate to PrP^C^ due to its higher affinity. However, while the combination spectrum preserves the superhyperfine splitting attributed to ^15^N-PrP^C^, the intensity is reduced. Moreover, there are subtle shifts in A_∥_ and g_∥_ with values distinct from either Cu^2+^-PrP^C^ or Cu^2+^ Aβ, which suggests that Cu^2+^ is in a new coordination environment with Cu^2+^ simultaneously coordinating to both Aβ30 and PrP^C^. To elucidate this new coordination environment and identify the coordinating atoms, we applied ESEEM and HYSCORE.

### ESEEM analysis of PrP^C^ and Aβ30

ESEEM provides important insight into the nuclei magnetically coupled to the Cu^2+^ center, revealing the nuclear Zeeman, hyperfine, and quadrupole interactions. One of the key advantages of ESEEM is its ability to differentiate between the nuclear spin environments of the isotopically labeled and unlabeled proteins. Since PrP^C^ is uniformly ^15^N-labeled (I = ½) its nuclear spin states will be distinct from those of the natural isotopic Aβ30 (I = 1), as shown in Figure 2A. Specifically, each coupled ^15^N exists in one of two nuclear spins states 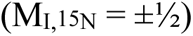 whereas quadrupolar ^14^N exists among three states, 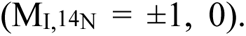 At X-band frequencies, the nuclear Zeeman and hyperfine terms of ^14^N in the higher energy manifold are in exact cancelation, thus selectively revealing the nuclear quadrupole transitions (19, 48, 49). Conversely, the nuclear hyperfine term in the ^15^N energy manifold results in two observable transitions, ν_α_ and ν_β_. These transitions are described by Equations 1-4, where A_iso_ is the isotropic hyperfine coupling, ν_nz_ is nuclear Zeeman frequency, T is the dipolar coupling, and θ is the angle between the electron-nuclear distance vector and the externally applied magnetic field (35, 50). By convention, we assign ν_α_ < ν_β_.

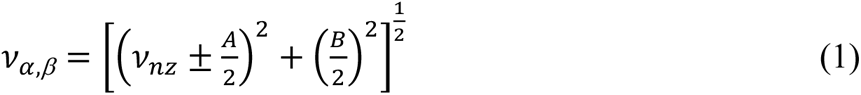

**Figure 2.**
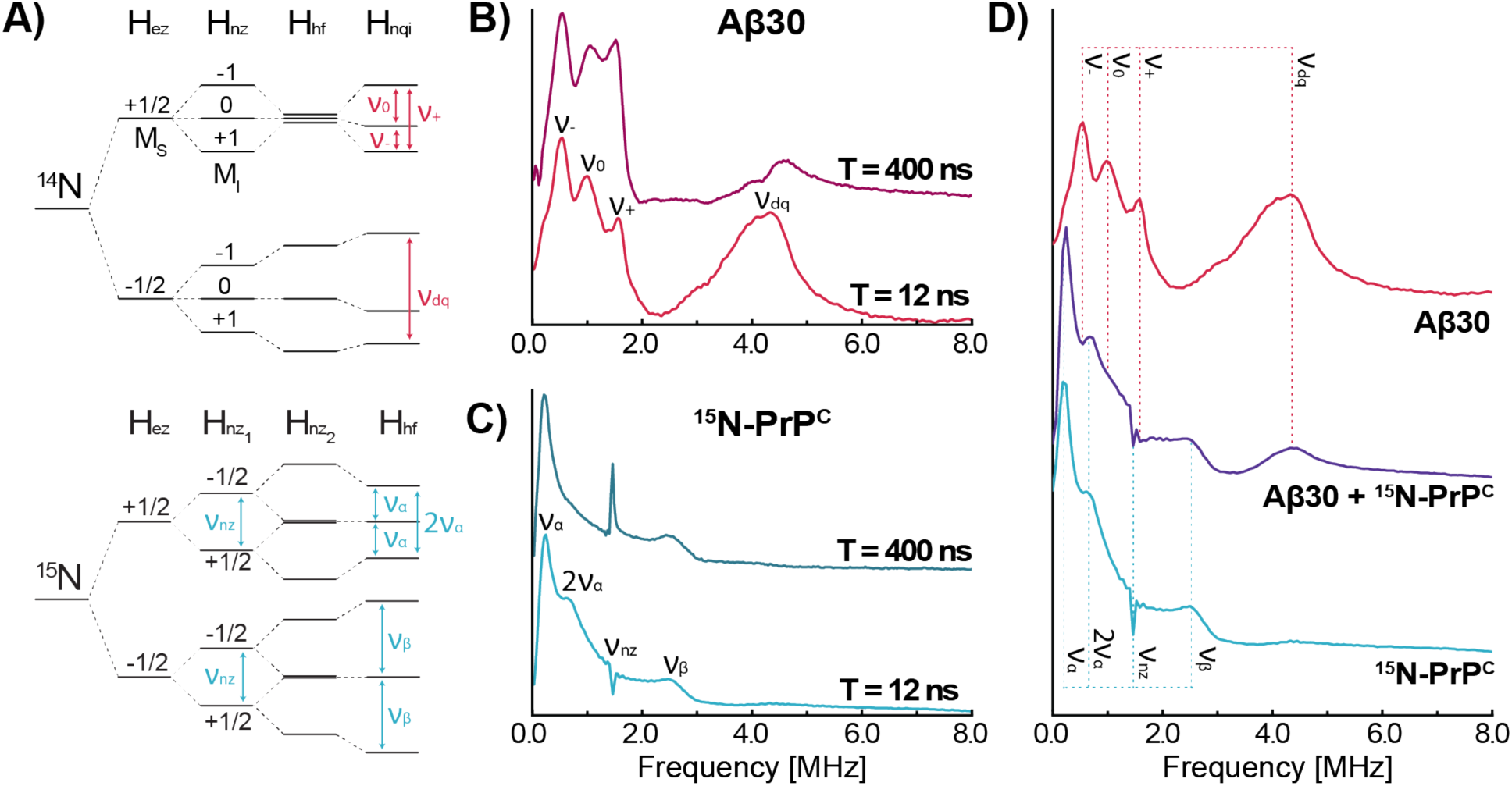
ESEEM transitions observed for ^14^N and ^15^N His coordination. A) The energy level diagrams for the Cu^2+^ electron interacting with either one ^14^N or two ^15^N are displayed, along with their relevant ESEEM transitions. B) ESEEM frequency domain of Aβ30 with Cu^2+^ measured with two different initial T values, 400 ns or 12 ns. Both spectra show the three clear NQI peaks labeled as ν_-_, ν_0_, ν_+_ and DQ peak labeled as ν_dq_. C) ESEEM frequency domain of ^15^N- with Cu^2+^, performed under the same conditions. There are three identifying peaks labeled as ν_*α*_, ν_β_, and ν_nz_. A fourth peak is observed at approximately 2ν_*α*_. D) The T = 12 ns traces for natural abundance Aβ30 (red, top), ^15^N-PrP^C^ (blue, bottom), and ^15^N-PrP^C^ + Aβ30 (purple, middle) with equimolar amounts of Cu^2+^.

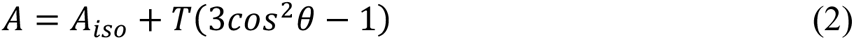

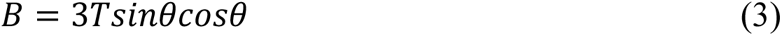

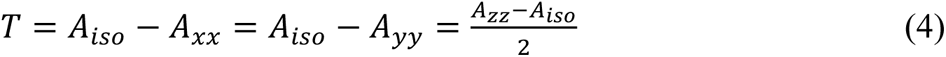

For histidine-Cu^2+^ coordination, relevant to our characterization here, ESEEM detects the noncoordinating nitrogen within the imidazole ring rather than the nitrogen directly coordinated to the paramagnetic center, outlined in Figure 1B,C (51).

The ESEEM spectra for Aβ30 and ^15^N-PrP^C^ are shown in Figure 2B and C, respectively. The spectrum of Aβ30 reveals two regions indicative of histidine-Cu^2+^ coupling (46, 52); the nuclear quadrupole interaction (NQI) peaks labeled as ν_-_ = 0.5 MHz, ν_0_ = 1.0 MHz, and ν_+_ = 1.5 MHz, along with the double quantum (DQ) peak, ν_dq_ = 4.2 MHz, consistent with previous work (18, 46, 52). Next, we observe four peaks in the ^15^N-PrP^C^ spectrum: ν_*α*_ = 0.3 MHz and ν_β_ = 2.5 MHz, consistent with previous work (53), ν_nz_ = 1.5 MHz, likely due to another weakly coupled ^15^N (i.e., small A_iso_) and a new peak at 0.6 MHz that has not been previously identified in other literature. Interestingly, this peak is at a frequency of double of the ν_*α*_ transition, which led us to investigate if this transition occurs when two or more equivalent ^15^N His are coupled to Cu^2+^. The ^15^N energy diagram (Figure 2A) shows that when two equivalent ^15^N His are coupled to the Cu^2+^ center, there is a potential 2ν_*α*_ transition. Using the program EasySpin (33), we simulated 3-pulsed ESEEM spectra with varying numbers of equivalent ^15^N His coupled to Cu^2+^, as shown in Figure S4. This analysis supports a 2ν_*α*_ transition occurring upon multiple equivalent ^15^N His coupling to Cu^2+^. Other possible sources for the 0.6 MHz transition, such as blind spots due to τ suppression and multiple nonequivalent ^15^N are explored in the subsequent section.

Before acquiring three-pulse ESEEM spectra of both ^15^N and ^14^N simultaneously, we set out to optimize pulse sequence parameters to provide the greatest intensity of all relevant frequency transitions. First, we examined how initial T changes the observed ESEEM spectra of Aβ30 and ^15^N-PrP^C^, as shown in Figure 2B and 2C respectively. In other published studies, a longer initial T values are used to allow adequate spacing between pulses (54). However, by decreasing initial T from 400 ns to 12 ns, as expected, we find that broad components assigned to multiple His coordination, notably the diagnostic 4.2 MHz DQ peak for ^14^N His and the 0.6 MHz for ^15^N, are properly resolved. Additionally, the shorter initial T does not compromise the three ^14^N NQI peaks. Therefore, we set the initial T to 12 ns for subsequent ESEEM as it is optimal for both ^14^N and ^15^N.

To optimize ESEEM acquisition with respect to τ, we developed a mathematical expression to incorporate two equivalent I = ½ nuclei in ESEEM from fundamental ESEEM equations, Equations S1-S11. Using Equation S11, we calculated the intensity of ν_*α*_, ν_β_, and 2ν_*α*_, as a function of τ, shown in Figure S5. From Figure S5, the optimal τ values for ν_*α*_ are between 140 – 250 ns. We therefore ran ESEEM experiments at τ = 144 ns and τ = 210 ns, noting that both have the added benefit of suppressing high-frequency proton ESEEM (24, 35). Figure S6, shows the comparison of τ = 144 ns and τ = 210 ns for both Aβ30 and ^15^N-PrP^C^. Notably, setting τ = 144 ns distorts the ^14^N peaks for Aβ30 and decreases the ν_β_ peak intensity for ^15^N-PrP^C^, as predicted from Equation S11 and shown in Figure S5. However, setting τ = 210 ns resolves all relevant transitions, including the diagnostic 2ν_α_. We therefore chose a τ = 210 ns for all subsequent ESEEM. The experimental ESEEM time domain signals are shown in Figure S7. Lastly, in optimizing T and τ, we did not focus on the ν_nz_ transition as it is not indicative of His coordination. Further analysis of this peak is discussed in the HYSCORE section.

In Figure 2D, we show the optimized ESEEM spectra for Aβ30, ^15^N-PrP^C^, each with one equivalent of Cu^2+^. The ESEEM of Cu^2+^ with both Aβ30 and ^15^N-PrP^C^, prepared in a 1:1:1 mixture, shows clearly the simultaneous features from both proteins, consistent with Cu^2+^ simultaneously coordinating to Aβ30 and PrP^C^. All ESEEM of Figure 2D are normalized to the echo intensity at T = 12 ns, therefore permitting analysis of the relative peak intensities among samples. With this normalization to echo intensity and, hence, concentration, we observe that the ^14^N ν_dq_ peak intensity in the 1:1:1 sample is persistent but decreased by 70% in the presence of equimolar ^15^N-PrP^C^. Noting that the intensity of the ν_dq_ transition is reflective of the number of coordinated ^14^N His imidazole (55), this suggests that Aβ His coordination to Cu^2+^ is reduced from two or three His (as reported for the Aβ-Cu^2+^ complex depending on pH and buffer conditions(16)), to one His upon the addition of ^15^N-PrP^C^. Furthermore, the 2ν_*α*_ peak, indicative of multiple ^15^N-PrP^C^ His coordination, remains pronounced in the 1:1:1 combination sample. As noted in the previous section, the dissociation constant of the PrP^C^-Cu^2+^ complex is approximately 30x lower than that of the Aβ-Cu^2+^ complex. As such, the ESEEM spectrum of the 1:1:1 sample should be dominated by the interaction between PrP^C^ and Cu^2+^. Observation of a persistent ^14^N signal in this mixture is therefore supportive of a ternary complex with both proteins simultaneously coordinating to the Cu^2+^ center. (Note: For reference, Figure S8 compares ^14^N and ^15^N histidine ESEEM at equal concentrations). This tentative assignment is further tested with 2D HYSCORE experiments.

### ^15^N/^14^N HYSCORE

The ^14^N and ^15^N signals from the ESEEM transitions below 1.5 MHz exhibit significant overlap – we therefore performed HYSCORE (hyperfine sublevel correlation) EPR, which is sensitive to the same coupled nuclear transitions as ESEEM, but introduces an additional π pulse and evolution period, thereby providing a two-dimensional spectrum with increased peak-to-peak resolution, along with correlations between nuclear spin manifolds (19, 24, 25).

The HYSCORE spectrum of ^15^N-PrP^C^-Cu^2+^ is shown in Figure 3A, revealing intense cross peaks symmetric across the diagonal at (0.32 MHz, 2.68 MHz) and (2.68 MHz, 0.32 MHz), corresponding to (ν_α_, ν_β_), and the diagonally symmetric (ν_β_, ν_α_), respectively. Similar spectra have been observed for Cu^2+^ coordination to glycine (19) and histidine (52), as well as an interaction between organic radicals and histidine (56, 57). Proximal to these intense peaks, we also observe weaker transitions at (0.63 MHz, 2.74 MHz) and (2.74 MHz, 0.63 MHz) consistent with (2ν_α_, ν_β_) cross peaks. Finally, a third peak is observed along the diagonal at (1.47 MHz, 1.47 MHz), which matches the ^15^N Larmor frequency. We associate the (1.47 MHz, 1.47 MHz) peak to weakly coupled ^15^N, as supported by Equations 1-3 in which ν_α_ and ν_β_ both converge to ν_nz_, in the limit of small A_iso_ and T. As shown in Figure S9, HYSCORE simulations in this limit predict a weak ν_nz_ peak along the diagonal when compared to calculated off diagonal peaks calculate for coupled ^15^N His. Furthermore, we observe that incorporating multiple weakly coupled ^15^N does not change the ν_nz_ peak, as seen in Figure S10. Thus, this diagonal peak is assigned to the collective contribution from the rich bath of weakly coupled ^15^N existing at longer distances from the Cu^2+^ center throughout uniformly labeled PrP^C^.

**Figure 3.**
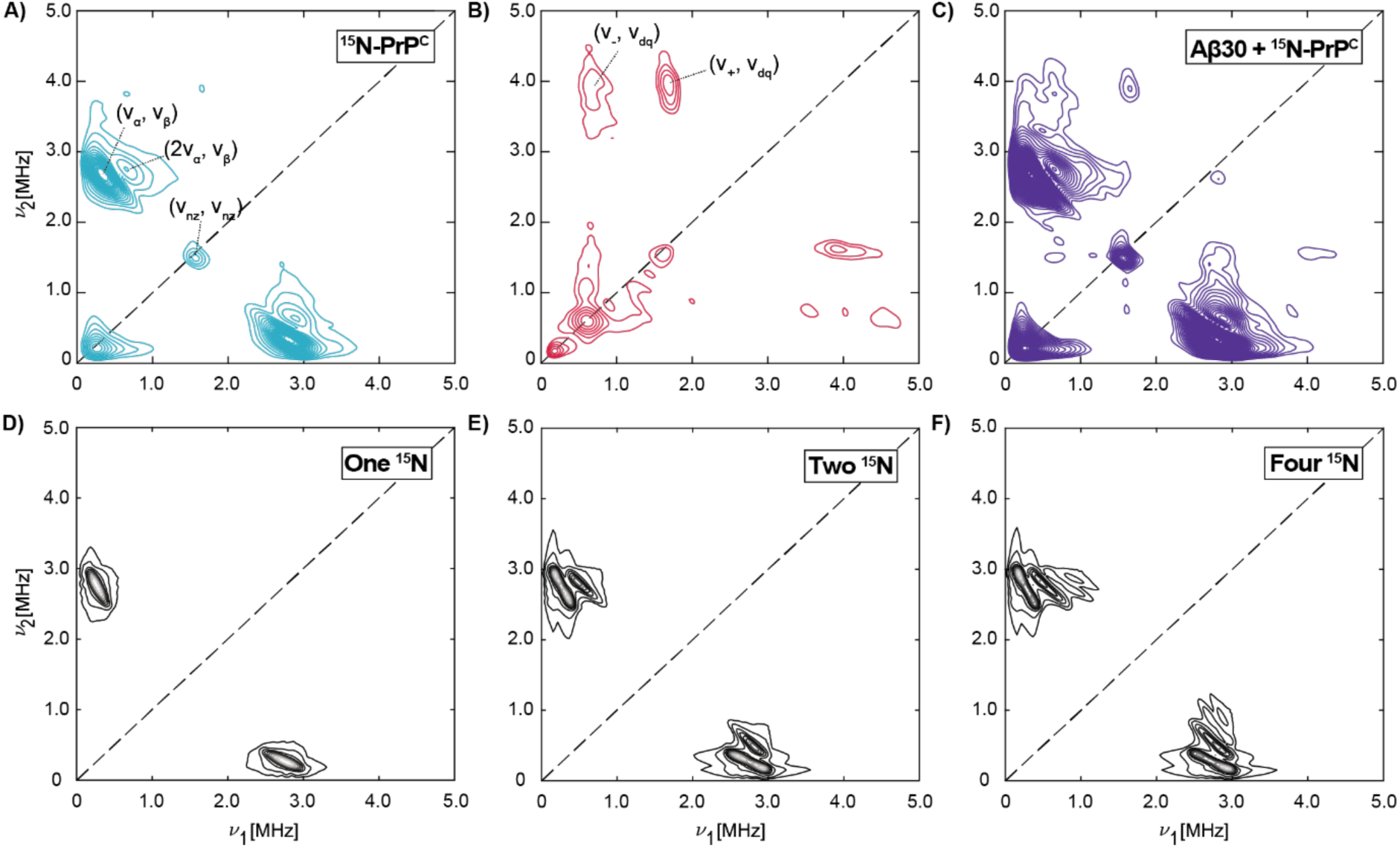
HYSCORE Contour plots for ^15^N and ^14^N His coordination. A) ^15^N-PrP^C^ (blue), B) natural abundance Aβ30 (red), and C) Aβ30 + ^15^N-PrP^C^ (purple). Spectra were acquired with 100 µM Cu^2+^, 100 µM Aβ30, and 100 µM ^15^N-PrP^C^. The black dashed line represents the equation ν_1_ = ν_2_. D) Simulated ^15^N HYSCORE spectra using one coupled ^15^N. τ was set to 210 ns to match the experimental τ. At this τ, only one peak is observed, implying no presence of a blind spot. Simulated HYSCORE with E) two and F) four coupled ^15^N nucleus were run with varied τ from 100 ns to 400 ns in 20 ns increments to account for any blind spots. Hyperfine values used to simulate these data are A_xx_ = A_yy_ = 3.2 MHz, A_zz_ = 2.0 MHz, A_iso_ = 2.8 MHz, and T = -0.4 MHz.

Next, we obtained HYSCORE spectra from natural abundance Aβ30, with and without ^15^N-PrP^C^, as shown in Figure 3B and C. These HYSCORE spectra were obtained with the same samples used in Figure 2. Focusing on natural abundance Aβ30, the reported HYSCORE contains cross peaks at (0.6 MHz, 3.8 MHz) and (1.6 MHz, 3.9 MHz), as well as similar peaks symmetric across the diagonal. These peaks are consistent with the (ν_-_, ν_dq_) and (ν_+_, ν_dq_) transitions observed when Cu^2+^ is coupled to the imidazole ring in histidine (19). Furthermore, the absence of a (ν_+_, ν_dq_) cross peak around (2.8 MHz, 4.0 MHz) indicates the absence of direct backbone amide coordination, consistent with current coordination models of Cu^2+^ to Aβ30 (10, 46). When both Aβ30 and ^15^N-PrP^C^ are present in the 1:1:1 mixture with copper (Figure 3C), we are able to clearly resolve the aforementioned peaks (ν_-_, ν_dq_), (ν_+_, ν_dq_) from ^14^N-His, and (ν_α_, ν_β_) from ^15^N His. Importantly, the (2ν_α_, ν_β_) cross peak persists, demonstrating multiple ^15^N-His from PrP^C^ coordinated to Cu^2+^ in the presence of Aβ30. Together, Figure 3C reveals clear, distinguishable, simultaneous His-Cu^2+^ signals from both Aβ30 and ^15^N-PrP^C^, with the latter contributing at least two His to the complex. Our interpretation of Cu^2+^ coordination to multiple His from PrP^C^ and at least one His from Aβ30 are reasoned on the grounds that the 2ν_α_ is due to Cu^2+^ coupling to two equivalent ^15^N. However, there are other possible sources for multiple peaks occurring in HYSCORE spectra. For example, selection of a specific τ value can result in blind spots, in which certain cross peaks are split, giving the appearance of two adjacent peaks (24, 33, 58). To explore this possibility, we performed a HYSCORE simulation for a single coupled ^15^N incorporating the same τ used in the experiment. As shown in Figure 3D, the simulation results in a single peak with no observable blind spot. This is further explored in Figure S11, which shows the results from several τ values. When ^15^N HYSCORE simulations do result in a blind spot, the observed cross peaks are not consistent with the (2ν_α_, ν_β_) cross peak observed experimentally.

Next, we performed HYSCORE simulations for two and four coupled ^15^N (Figure 3E and 3F), using the same parameters from Figure 3D. These simulations show that incorporation of equivalent ^15^N produces a progression of new, higher frequency peaks at multiples of ν_α_ and fixed ν_β_. Specifically, adjacent to the expected (ν_α_, ν_β_) is a cross peak at (2ν_α_, ν_β_), consistent with experimental spectra. This is in contrast to the observation of only a single cross peak reported in previous work done on systems with a single coupled ^15^N (19, 52, 57). Focusing next on the relative intensity of the observed cross peaks, the simulations find an intensity ratio (measured by peak height) of 1.0:0.6 between the original (ν_α_, ν_β_) and the (2ν_α_, ν_β_), which is higher than 1.0:0.3 observed experimentally. However, the ratios depend strongly on the broadening and the use of τ blind spot suppression in the simulations as well as the apodization used when processing the experimental HYSCORE data. Moving to four nuclei, the simulated HYSCORE contains a set of four peaks with an intensity ratio of 1.0:0.7:0.1:0.1. Because of the relatively low intensity of the latter two peaks, it would be difficult to observe these experimentally. Thus, it is feasible to differentiate between single and multiple ^15^N but determining the exact number of coupled ^15^N remains challenging through this analysis. Nevertheless, the assignment of at least two coupled ^15^N nuclei is clear in both the ^15^N-PrP-Cu^2+^ complex and the 1:1:1 Aβ, PrP^C^, Cu^2+^ mixture.

HYSCORE simulations in Figure S12, show that the (2ν_α_, ν_β_) cross peak is resolved only when the magnitude of the dipolar coupling, T, is on the order of -0.4 MHz or greater. Simulations with a lower magnitude dipolar coupling, T = -0.1 MHz reveal solely the (ν_α_, ν_β_) cross peak. Moreover, at T = -1.2 MHz, a third cross peak appears at (2ν_α_, 2ν_β_), Figure S13. Similar simulations with constant T but varying A_iso_ show little influence on the presence of (2ν_α_, ν_β_) or (2ν_α_, 2ν_β_) (Figure S14). Based on the analyses above, we assign the (2ν_α_, ν_β_) peak observed in both ESEEM and HYSCORE to the ^15^N-His ΔI = ±2 transition, outlined in Figure 2A.

## Conclusions

The aggregation of IDPs, frequently stabilized by metals and other proteins, is a key factor in the onset and progression of neurodegenerative diseases. While it is well established that PrP^C^ serves as a major cellular receptor for Aβ and plays a crucial role in both Aβ translocation into cells and progression of the disease, the molecular mechanisms underlying the relevant intermolecular contacts are unknown. We hypothesize that the bimolecular interaction between Aβ and PrP^C^ is stabilized or influenced by Cu^2+^. This is supported by the fact that both proteins have nM affinity for Cu^2+^, as well as Cu^2+^ being highly concentrated in amyloid plaques. In this study, we employed various spectroscopic techniques, including CW-EPR, ESEEM, and HYSCORE, to observe and characterize Cu^2+^ coordination to both proteins. Our findings provide spectral evidence indicative of Cu^2+^ coordinating simultaneously to both Aβ30 and PrP^C^. By CW-EPR, we identified that this coordination involves histidine residues from both proteins. By ESEEM and HYSCORE, we identified that when both proteins are present, there is a contribution from ^14^N and ^15^N histidine. Furthermore, we observed that upon addition of PrP^C^ to Aβ30-Cu^2+^, the ^14^N-His ν_dq_ peak intensity decreases by 70%, suggesting a reduced His contribution from Aβ30. However, His coordination from Aβ persists despite its significantly lower affinity compared to that for the PrP^C^-Cu^2+^ complex. The (2ν_α_, ν_β_) assigned to two or more His from ^15^N-PrP^C^ remains in the presence of Aβ. Taken together, our data suggest that three His from PrP^C^ and a single His from Aβ30 coordinate to Cu^2+^ in the formation of a ternary complex. Our findings are summarized in Figure 4.

**Figure 4.**
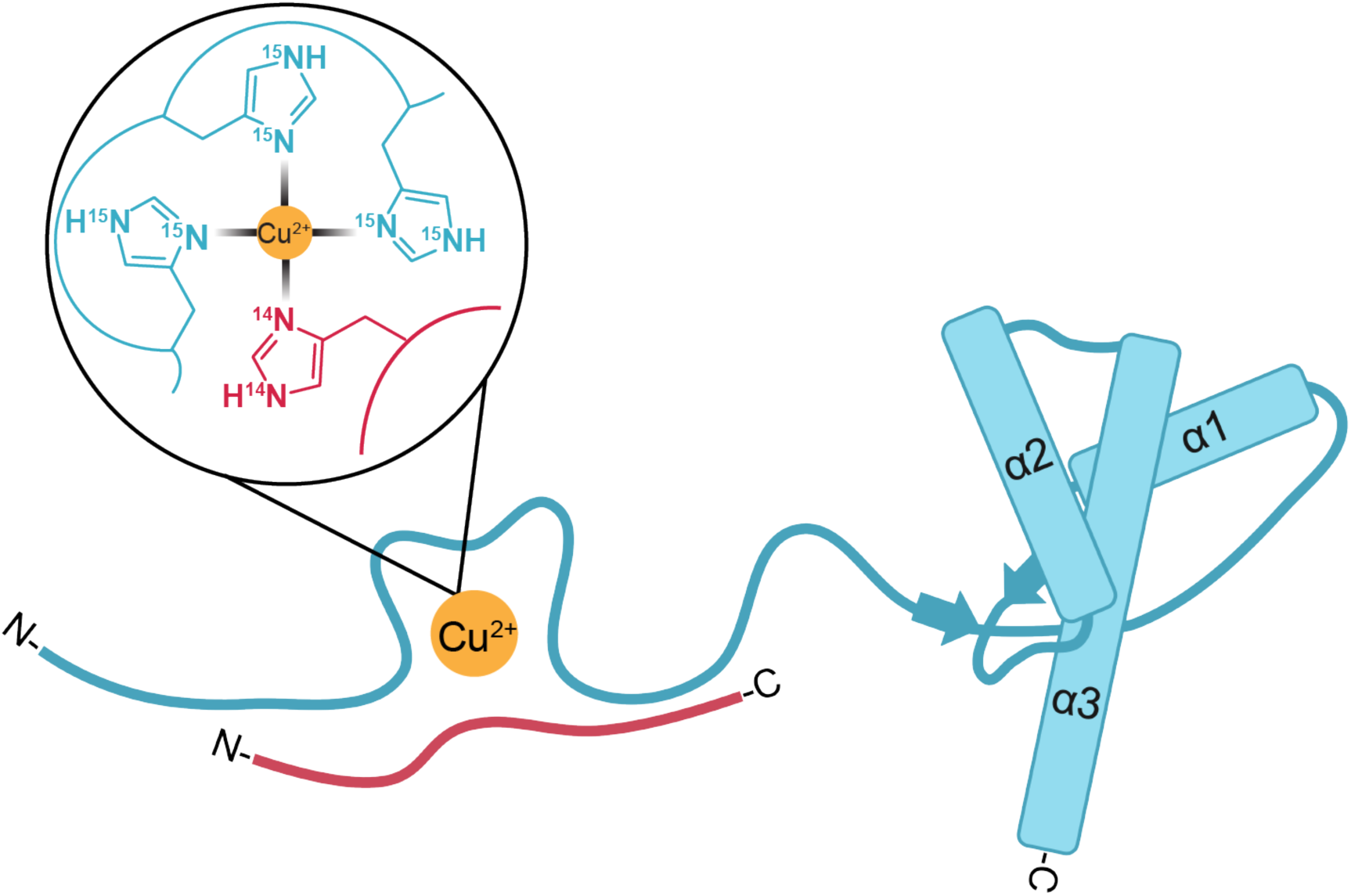
Structural model of natural abundance Aβ and ^15^N-PrP^C^ coordinating to Cu^2+^. The schematic structure of PrP^C^ is shown in blue, identifying its N-terminal IDP domain and its structured C-terminal domain, defined by three α-helices and one anti-parallel β-sheet. Our findings suggest that the disordered N-terminal domain coordinates Cu^2+^ through three ^15^N-His residues in the OR domain. Aβ is represented as an IDP (red) coordinating Cu^2+^ through a single ^14^N-His. The coordination of the three ^15^N-His and one ^14^N-His to Cu^2+^ is shown in the insert.

In addition to characterization of this novel complex, we also developed a method that identifies multiple ^15^N-His coordination to copper centers and demonstrates the feasibility and utility of ^14^N and ^15^N mixed isotope experiments, especially as they apply to neurodegenerative proteins. This approach is readily adapted to segmentally labeled proteins, for example, to identify key residues or segments involved in metal regulation or chelation. Importantly, this study demonstrates that Aβ, PrP^C^, and Cu^2+^ interact and form a ternary complex at physiological pH, suggesting a potential mechanism for cellular uptake of Aβ, a crucial step in the progression of AD.

## Associated Content

Supporting information.

## Author Contributions

**Amanda Smart**: Conceptualization (Lead), Methodology (Equal), Validation (Lead), Formal Analysis (Equal), Investigation (Lead), Resources (Supporting), Data Curation (Equal), Writing – Original draft (Equal), Writing – Review & Editing (Equal), Visualization (Supporting), and Project Administration (Equal). **Kevin Singewald**: Conceptualization (Supporting), Methodology (Equal), Software (Lead), Validation (Supporting), Formal Analysis (Equal), Investigation (Supporting), Resources (Supporting), Data Curation (Equal), Writing – Original draft (Equal), Writing – Review & Editing (Equal), Visualization (Lead), Supervision (Supporting), Project Administration (Equal), and Funding acquisition (Supporting). **Zikri Hasanbasri**: Methodology (Supporting) and Writing – Review & Editing (Supporting). **R. David Britt**: Writing – Review & Editing (Supporting), Supervision (Supporting) and Funding acquisition (Supporting). **Glenn L. Millhauser**: Conceptualization (Lead), Validation (Supporting), Resources (Lead), Writing – Original draft (Equal), Writing – Review & Editing (Equal), Supervision (Lead), Project Administration (Equal), and Funding acquisition (Lead).

## Acknowledgement

This research was supported by the National Institutes of Health (R35GM131781, A.S., K.S., G.L.M), the Cinvestav and UC Alianza MX (A.S., K.S., Z.H., R.D.B, G.L.M.), National Institute of General Medicine Sciences Institutional Research and Academic Career Development Award (K12GM139185, K.S.), and the Initiative for Maximizing Student Development Program, IMSD (1T32GM135742-02, A.S.). The pulsed EPR instrument was funded through NIH grant S10OD024980. John L. McCracken is thanked for helpful discussion on ESEEM and HYSCORE results.

## Abbreviations

Aβ30: amyloid-β residues 1-30
AD: Alzheimer’s disease
CW: continuous wave
DQ: double quantum
EPR: electron paramagnetic resonance
ESEEM: electron spin echo envelope modulation
HYSCORE: hyperfine sublevel correlation
IDP: intrinsically disordered peptide
NQI: nuclear quadrupole interaction
OR: octapeptide repeat
PrP^C^: cellular isoform of prion protein residues 23-230

## REFERENCES

1. O’Brien, R. J., and Wong, P. C. (2011) Amyloid Precursor Protein Processing and Alzheimer’s Disease. Annu. Rev. Neurosci. 34, 185–204

2. Laurén, J., Gimbel, D. A., Nygaard, H. B., Gilbert, J. W., and Strittmatter, S. M. (2009) Cellular prion protein mediates impairment of synaptic plasticity by amyloid-β oligomers. Nature. 457, 1128–1132

3. Schwarze-Eicker, K., Keyvani, K., Görtz, N., Westaway, D., Sachser, N., and Paulus, W. (2005) Prion protein (PrPc) promotes β-amyloid plaque formation. Neurobiology of Aging. 26, 1177–1182

4. Foley, A. R., Roseman, G. P., Chan, K., Smart, A., Finn, T. S., Yang, K., Lokey, R. S., Millhauser, G. L., and Raskatov, J. A. (2020) Evidence for aggregation-independent, PrP ^C^ - mediated Aβ cellular internalization. Proc Natl Acad Sci USA. 117, 28625–28631

5. Smith, L. M., Kostylev, M. A., Lee, S., and Strittmatter, S. M. (2019) Systematic and standardized comparison of reported amyloid-β receptors for sufficiency, affinity, and Alzheimer’s disease relevance. Journal of Biological Chemistry. 294, 6042–6053

6. Legname, G., and Scialò, C. (2020) On the role of the cellular prion protein in the uptake and signaling of pathological aggregates in neurodegenerative diseases. Prion. 14, 257–270

7. LaFerla, F. M., Green, K. N., and Oddo, S. (2007) Intracellular amyloid-β in Alzheimer’s disease. Nat Rev Neurosci. 8, 499–509

8. Haas, L. T., Kostylev, M. A., and Strittmatter, S. M. (2014) Therapeutic Molecules and Endogenous Ligands Regulate the Interaction between Brain Cellular Prion Protein (PrPC) and Metabotropic Glutamate Receptor 5 (mGluR5). Journal of Biological Chemistry. 289, 28460–28477

9. Larson, M., Sherman, M. A., Amar, F., Nuvolone, M., Schneider, J. A., Bennett, D. A., Aguzzi, A., and Lesné, S. E. (2012) The Complex PrP ^c^ -Fyn Couples Human Oligomeric Aβ with Pathological Tau Changes in Alzheimer’s Disease. J. Neurosci. 32, 16857–16871

10. Faller, P., and Hureau, C. (2009) Bioinorganic chemistry of copper and zinc ions coordinated to amyloid-β peptide. Dalton Trans. 10.1039/B813398K

11. Miller, L. M., Wang, Q., Telivala, T. P., Smith, R. J., Lanzirotti, A., and Miklossy, J. (2006) Synchrotron-based infrared and X-ray imaging shows focalized accumulation of Cu and Zn co-localized with β-amyloid deposits in Alzheimer’s disease. Journal of Structural Biology. 155, 30–37

12. Atwood, C. S., Moir, R. D., Huang, X., Scarpa, R. C., Bacarra, N. M. E., Romano, D. M., Hartshorn, M. A., Tanzi, R. E., and Bush, A. I. (1998) Dramatic Aggregation of Alzheimer Aβ by Cu(II) Is Induced by Conditions Representing Physiological Acidosis. Journal of Biological Chemistry. 273, 12817–12826

13. Reddy, P. H., and Beal, M. F. (2008) Amyloid beta, mitochondrial dysfunction and synaptic damage: implications for cognitive decline in aging and Alzheimer’s disease. Trends in Molecular Medicine. 14, 45–53

14. Tassone, G., Kola, A., Valensin, D., and Pozzi, C. (2021) Dynamic Interplay between Copper Toxicity and Mitochondrial Dysfunction in Alzheimer’s Disease. Life. 11, 386

15. Karr, J. W., and Szalai, V. A. (2008) Cu(II) Binding to Monomeric, Oligomeric, and Fibrillar Forms of the Alzheimer’s Disease Amyloid-β Peptide. Biochemistry. 47, 5006–5016

16. Faller, P. (2009) Copper and Zinc Binding to Amyloid-β: Coordination, Dynamics, Aggregation, Reactivity and Metal-Ion Transfer. ChemBioChem. 10, 2837–2845

17. Walter, E. D., Chattopadhyay, M., and Millhauser, G. L. (2006) The Affinity of Copper Binding to the Prion Protein Octarepeat Domain: Evidence for Negative Cooperativity. Biochemistry. 45, 13083–13092

18. Aronoff-Spencer, E., Burns, C. S., Avdievich, N. I., Gerfen, G. J., Peisach, J., Antholine, W. E., Ball, H. L., Cohen, F. E., Prusiner, S. B., and Millhauser, G. L. (2000) Identification of the Cu ^2+^ Binding Sites in the N-Terminal Domain of the Prion Protein by EPR and CD Spectroscopy. Biochemistry. 39, 13760–13771

19. Burns, C. S., Aronoff-Spencer, E., Legname, G., Prusiner, S. B., Antholine, W. E., Gerfen, G. J., Peisach, J., and Millhauser, G. L. (2003) Copper Coordination in the Full-Length, Recombinant Prion Protein. Biochemistry. 42, 6794–6803

20. Rudd, P. (2002) Glycosylation and prion protein. Current Opinion in Structural Biology. 12, 578–586

21. Schilling, K. M., Tao, L., Wu, B., Kiblen, J. T. M., Ubilla-Rodriguez, N. C., Pushie, M. J., Britt, R. D., Roseman, G. P., Harris, D. A., and Millhauser, G. L. (2020) Both N-Terminal and C-Terminal Histidine Residues of the Prion Protein Are Essential for Copper Coordination and Neuroprotective Self-Regulation. Journal of Molecular Biology. 432, 4408–4425

22. Schilling, K. M., Jorwal, P., Ubilla-Rodriguez, N. C., Assafa, T. E., Gatdula, J. R. P., Vultaggio, J. S., Harris, D. A., and Millhauser, G. L. (2023) N-glycosylation is a potent regulator of prion protein neurotoxicity. Journal of Biological Chemistry. 299, 105101

23. Wu, B., McDonald, A. J., Markham, K., Rich, C. B., McHugh, K. P., Tatzelt, J., Colby, D. W., Millhauser, G. L., and Harris, D. A. (2017) The N-terminus of the prion protein is a toxic effector regulated by the C-terminus. eLife. 6, e23473

24. Dikanov, S. A., and Tsvetkov, Y. D. (1992) Electron spin echo envelope modulation (ESEEM) spectroscopy, CRC Press, Boca Raton

25. Höfer, P., Grupp, A., Nebenführ, H., and Mehring, M. (1986) Hyperfine sublevel correlation (hyscore) spectroscopy: a 2D ESR investigation of the squaric acid radical. Chemical Physics Letters. 132, 279–282

26. Deligiannakis, Y., Jegerschöld, C., and William Rutherford, A. (1997) EPR and ESEEM study of the plastoquinone anion radical QA−. in photosystem II treated at high pH. Chemical Physics Letters. 270, 564–572

27. Flanagan, H. L., Gerfen, G. J., Lai, A., and Singel, D. J. (1988) Orientation-selective 14N electron spin echo envelope modulation (ESEEM): The determination of 14N quadrupole coupling tensor principal axis orientations in orientationally disordered solids. The Journal of Chemical Physics. 88, 2162–2168

28. Deligiannakis, Y., Louloudi, M., and Hadjiliadis, N. (2000) Electron spin echo envelope modulation (ESEEM) spectroscopy as a tool to investigate the coordination environment of metal centers. Coordination Chemistry Reviews. 204, 1–112

29. Evans, E. G. B., Pushie, M. J., Markham, K. A., Lee, H.-W., and Millhauser, G. L. (2016) Interaction between Prion Protein’s Copper-Bound Octarepeat Domain and a Charged C-Terminal Pocket Suggests a Mechanism for N-Terminal Regulation. Structure. 24, 1057– 1067

30. Spevacek, A. R., Evans, E. G. B., Miller, J. L., Meyer, H. C., Pelton, J. G., and Millhauser, G. L. (2013) Zinc Drives a Tertiary Fold in the Prion Protein with Familial Disease Mutation Sites at the Interface. Structure. 21, 236–246

31. Gibson, D. G., Young, L., Chuang, R.-Y., Venter, J. C., Hutchison, C. A., and Smith, H. O. (2009) Enzymatic assembly of DNA molecules up to several hundred kilobases. Nat Methods. 6, 343–345

32. Fielding, A. J., Fox, S., Millhauser, G. L., Chattopadhyay, M., Kroneck, P. M. H., Fritz, G., Eaton, G. R., and Eaton, S. S. (2006) Electron spin relaxation of copper(II) complexes in glassy solution between 10 and 120K. Journal of Magnetic Resonance. 179, 92–104

33. Stoll, S., and Schweiger, A. (2006) EasySpin, a comprehensive software package for spectral simulation and analysis in EPR. Journal of Magnetic Resonance. 178, 42–55

34. Casto, J., Mandato, A., and Saxena, S. (2021) dHis-troying Barriers: Deuteration Provides a Pathway to Increase Sensitivity and Accessible Distances for Cu ^2+^ Labels. J. Phys. Chem. Lett. 12, 4681–4685

35. Stoll, S., Calle, C., Mitrikas, G., and Schweiger, A. (2005) Peak suppression in ESEEM spectra of multinuclear spin systems. Journal of Magnetic Resonance. 177, 93–101

36. Fábregas Ibáñez, L., Soetbeer, J., Klose, D., Tinzl, M., Hilvert, D., and Jeschke, G. (2019) Non-uniform HYSCORE: Measurement, processing and analysis with Hyscorean. Journal of Magnetic Resonance. 307, 106576

37. The Mathworks Inc. MATLAB version: 9.13

38. Thakur, A. K., Srivastava, A. K., Srinivas, V., Chary, K. V. R., and Rao, C. M. (2011) Copper Alters Aggregation Behavior of Prion Protein and Induces Novel Interactions between Its N- and C-terminal Regions. Journal of Biological Chemistry. 286, 38533–38545

39. You, H., Tsutsui, S., Hameed, S., Kannanayakal, T. J., Chen, L., Xia, P., Engbers, J. D. T., Lipton, S. A., Stys, P. K., and Zamponi, G. W. (2012) Aβ neurotoxicity depends on interactions between copper ions, prion protein, and *N* -methyl-D -aspartate receptors. Proc. Natl. Acad. Sci. U.S.A. 109, 1737–1742

40. Khosravani, H., Zhang, Y., Tsutsui, S., Hameed, S., Altier, C., Hamid, J., Chen, L., Villemaire, M., Ali, Z., Jirik, F. R., and Zamponi, G. W. (2008) Prion protein attenuates excitotoxicity by inhibiting NMDA receptors. J Gen Physiol. 131, i5–i5

41. Salzano, G., Giachin, G., and Legname, G. (2019) Structural Consequences of Copper Binding to the Prion Protein. Cells. 8, 770

42. Chen, S., Yadav, S. P., and Surewicz, W. K. (2010) Interaction between Human Prion Protein and Amyloid-β (Aβ) Oligomers. Journal of Biological Chemistry. 285, 26377–26383

43. Fluharty, B. R., Biasini, E., Stravalaci, M., Sclip, A., Diomede, L., Balducci, C., La Vitola, P., Messa, M., Colombo, L., Forloni, G., Borsello, T., Gobbi, M., and Harris, D. A. (2013) An N-terminal Fragment of the Prion Protein Binds to Amyloid-β Oligomers and Inhibits Their Neurotoxicity in Vivo. Journal of Biological Chemistry. 288, 7857–7866

44. Jones, C. E., Klewpatinond, M., Abdelraheim, S. R., Brown, D. R., and Viles, J. H. (2005) Probing Copper2+ Binding to the Prion Protein Using Diamagnetic Nickel2+ and 1H NMR: The Unstructured N terminus Facilitates the Coordination of Six Copper2+ Ions at Physiological Concentrations. Journal of Molecular Biology. 346, 1393–1407

45. Quintanar, L., and Millhauser, G. L. (2022) EPR of copper centers in the prion protein. in Methods in Enzymology, pp. 297–314, Elsevier, 666, 297–314

46. Silva, K. I., Michael, B. C., Geib, S. J., and Saxena, S. (2014) ESEEM Analysis of Multi-Histidine Cu(II)-Coordination in Model Complexes, Peptides, and Amyloid-β. J. Phys. Chem. B. 118, 8935–8944

47. Posadas, Y., Parra-Ojeda, L., Perez-Cruz, C., and Quintanar, L. (2021) Amyloid β Perturbs Cu(II) Binding to the Prion Protein in a Site-Specific Manner: Insights into Its Potential Neurotoxic Mechanisms. Inorg. Chem. 60, 8958–8972

48. Lee, H.-I., Doan, P. E., and Hoffman, B. M. (1999) General Analysis of 14N (I = 1) Electron Spin Echo Envelope Modulation. Journal of Magnetic Resonance. 140, 91–107

49. Peisach, J., and Blumberg, W. E. (1974) Structural implications derived from the analysis of electron paramagnetic resonance spectra of natural and artificial copper proteins. Archives of Biochemistry and Biophysics. 165, 691–708

50. Lin, C. P., Bowman, M. K., and Norris, J. R. (1986) Analysis of electron spin echo modulation in randomly oriented solids: 15N modulation of radical cations of bacteriochlorophyll *a* and the primary donor of photosynthetic bacterium Rhodospirillumbrum. The Journal of Chemical Physics. 85, 56–62

51. McCracken, J., Peisach, J., and Dooley, D. M. (1987) Cu(II) coordination chemistry of amine oxidases. Pulsed EPR studies of histidine imidazole, water, and exogenous ligand coordination. J. Am. Chem. Soc. 109, 4064–4072

52. Shin, B., and Saxena, S. (2008) Direct Evidence That All Three Histidine Residues Coordinate to Cu(II) in Amyloid-β _1−16_. Biochemistry. 47, 9117–9123

53. Dikanov, S., Felli, I., Viezzoli, M.-S., Spoyalov, A., and Hüttermann, J. (1994) X-band ESEEM spectroscopy of ^15^ N substituted native and inhibitor-bound superoxide dismutase: Hyperfine couplings with remote nitrogen of histidine ligands. FEBS Letters. 345, 55–60

54. Singewald, K., Wilkinson, J., and Saxena, S. (2021) Copper Based Site-directed Spin Labeling of Proteins for Use in Pulsed and Continuous Wave EPR Spectroscopy. BIO-PROTOCOL. 10.21769/BioProtoc.4258

55. McCracken, John., Pember, Stephen., Benkovic, S. J., Villafranca, J. J., Miller, R. J., and Peisach, Jack. (1988) Electron spin-echo studies of the copper binding site in phenylalanine hydroxylase from Chromobacterium violaceum. J. Am. Chem. Soc. 110, 1069–1074

56. Taguchi, A. T., O’Malley, P. J., Wraight, C. A., and Dikanov, S. A. (2014) Hyperfine and Nuclear Quadrupole Tensors of Nitrogen Donors in the Q _A_ Site of Bacterial Reaction Centers: Correlation of the Histidine N _δ_ Tensors with Hydrogen Bond Strength. J. Phys. Chem. B. 118, 9225–9237

57. Seif Eddine, M., Biaso, F., Arias-Cartin, R., Pilet, E., Rendon, J., Lyubenova, S., Seduk, F., Guigliarelli, B., Magalon, A., and Grimaldi, S. (2017) Probing the Menasemiquinone Binding Mode to Nitrate Reductase A by Selective ^2^ H and ^15^ N Labeling, HYSCORE Spectroscopy, and DFT Modeling. ChemPhysChem. 18, 2704–2714

58. Cox, N., Nalepa, A., Pandelia, M.-E., Lubitz, W., and Savitsky, A. (2015) Pulse Double-Resonance EPR Techniques for the Study of Metallobiomolecules. in Methods in Enzymology, pp. 211–249, Elsevier, 563, 211–249

