## Supporting Information for "Identifying the Copper Coordination Environment Between Interacting Neurodegenerative Proteins: A New Approach Using Pulsed EPR with ^14^N/^15^N Isotopic Labelling"

Amanda Smart, Kevin Singewald, Zikri Hasanbasri, R. David Britt, Glenn L. Millhauser

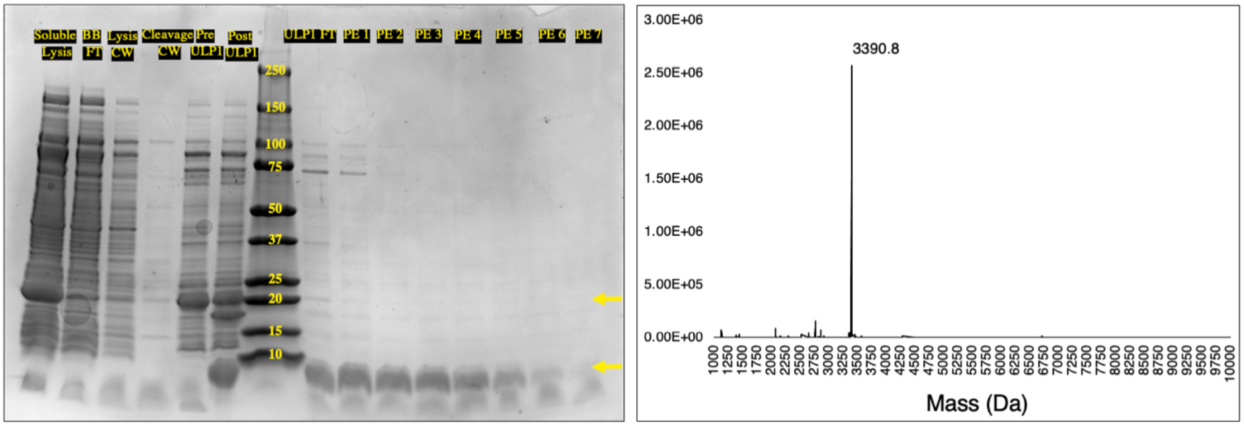

**Figure S1. SDS-PAGE and Mass Spectrometry confirmation of Aβ30 production.** SDS-PAGE (left) and mass spectrum (right) confirm the expression and purification of Aβ30. The gel shows lanes as stated from left to right: soluble cell lysate, $Ni^{2+}$-IMAC batch binding flow through, lysis buffer column wash, cleavage buffer column wash, pre-addition of ULP1 SUMO-protease, post-addition of ULP1 SUMO-protease, ULP1-cleave reaction flow through, protein elution buffer fraction 1-7. The cleavage of SUMO-Aβ30 is seen by the appearance of a band at 17.1 kDa for SUMO and 3.4 kDa for Aβ30 in the Post-ULP1 lane. The mass spectrum of purified Aβ30 shows that correct mass of 3390.8 Da.

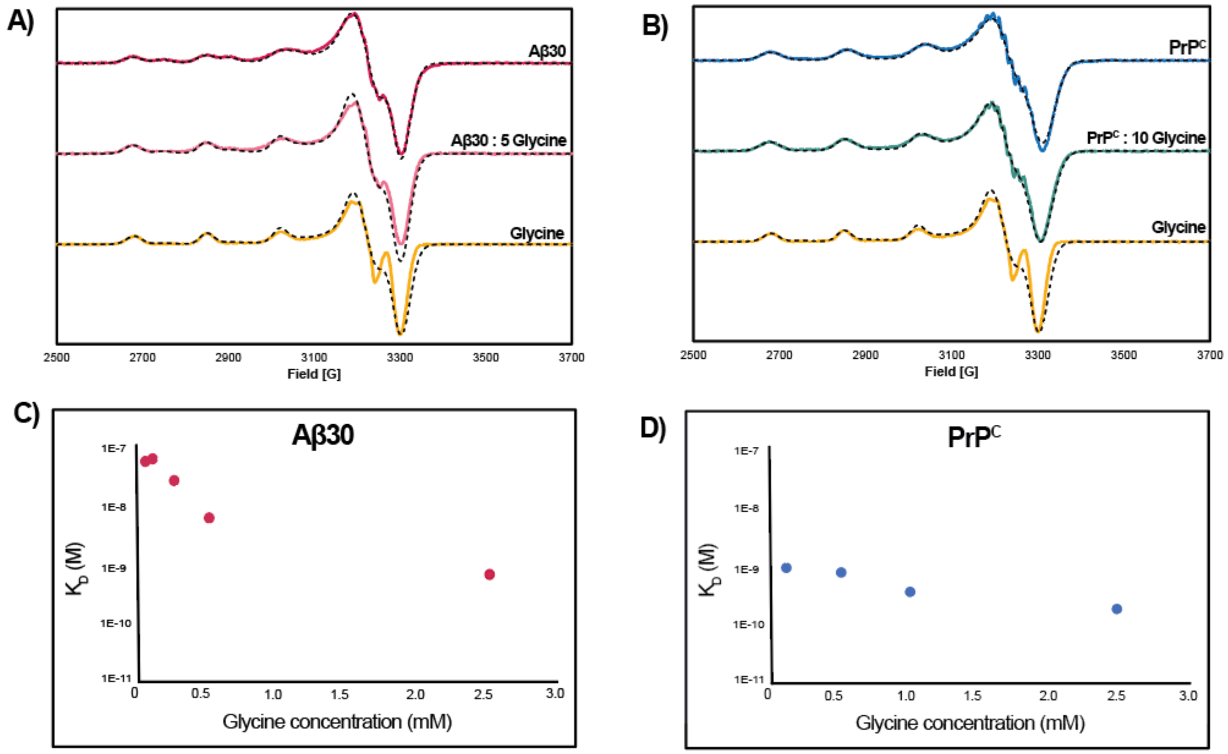

**Figure S2: Glycine competition assay to determine $Cu^{2+}$- protein binding affinities.** CW-EPR was used to calculate $K_D$ for Aβ30-$Cu^{2+}$ and $PrP^C$ -$Cu^{2+}$ in 50 mM HEPES, pH 7.4. Sample spectra for (A) Aβ30 and (B) $PrP^C$ in competition with glycine: (top) spectrum are of protein-$Cu^{2+}$ complex, (middle) is protein : X glycine : $Cu^{2+}$ complex, and (bottom) is glycine- $Cu^{2+}$ complex. Each were recorded with 100 µM protein, 100 µM $Cu^{2+}$, and 2500 µM glycine, except for in the protein glycine sample where glycine was 500 µM or 1000 µM, for Aβ30 or $PrP^C$, respectively. The dashed spectra overlaid are the EasySpin simulated spectra. The calculated $K_D$ for each experimental condition are plotted for Aβ30 (C) and $PrP^C$ (D). The calculated $K_D$ for Aβ30 and $PrP^C$ were 16.0 ± 14 nM and 0.519 ± 0.32 nM, respectively.

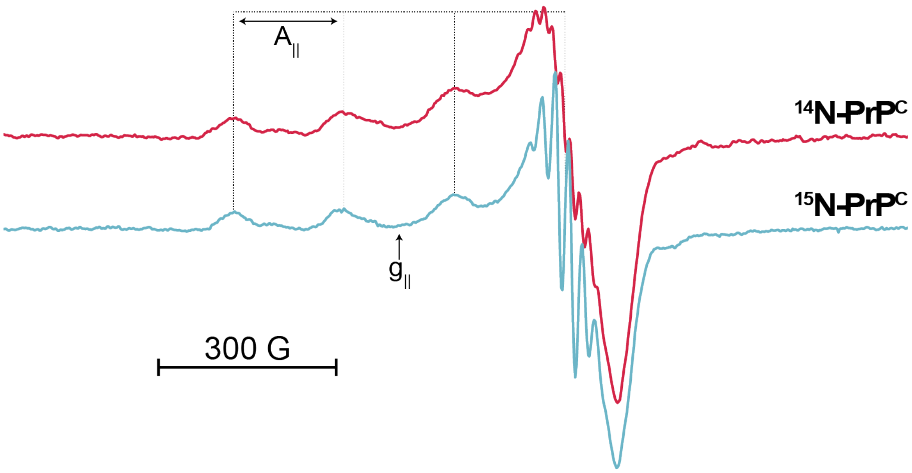

**Figure S3. Comparison between $^{14}N$ and $^{15}N$ CW-EPR.** Comparison between natural abundance $^{14}N$-$PrP^C$ (red) and $^{15}N$-$PrP^C$ (blue) CW-EPR. Both spectra have similar $A_{||}$ and $g_{||}$ values. One difference in the spectra is the superhyperfine observed in the high field perpendicular region. Because of the difference in nuclear spin, $I_{14N} = 1$ vs. $I_{15N} = ½$, the number of superhyperfine transitions changes. For four histidine coordination, $^{14}N$ would give a much broader signal with 9 superhyperfine transitions, whereas $^{15}N$ only has 5 transitions.

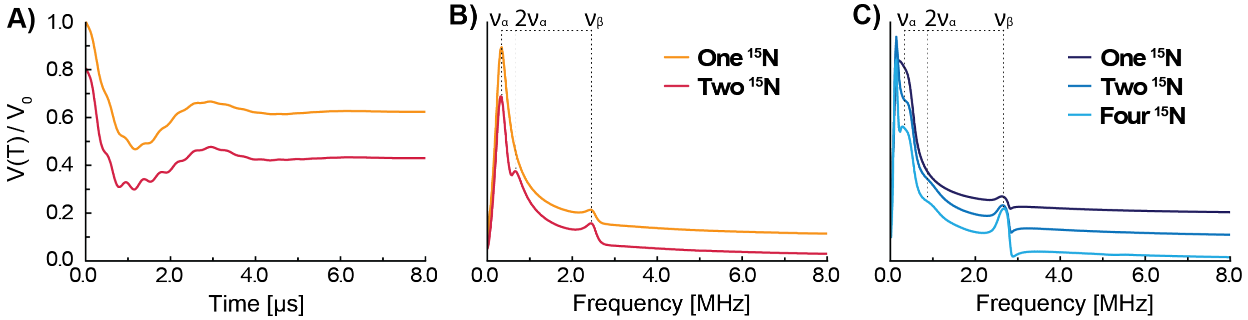

**Figure S4. 3-pulse ESEEM spectra calculated from fundamental ESEEM equations and simulated in EasySpin.** A) Calculated 3-pulse ESEEM time domain signal and B) respective Fourier transform of one (orange), and two (red) $^{15}N$ from Equation S8 and S11. $\nu_\alpha = 0.3$ MHz, $\nu_\beta = 2.5\ MHz$, $k = 0.2$, $\tau = 0.210\ \mu s$, and multiplying the time domain signal by $e^{-0.8T}$ to incorporate line broadening. The resulting spectra reports similar features to the spectra observed in Figures 2D, S6, and S7. C) EasySpin simulated ESEEM spectra of one, two, and four $^{15}N$. $A_{xx} = A_{yy} = 2.9$ MHz, $A_{zz} = 1.7$ MHz, $A_{iso} = 2.5$ MHz, T = -0.4 MHz. Beyond a single $^{15}N$, an additional peak appears around twice that of $\nu_\alpha$.

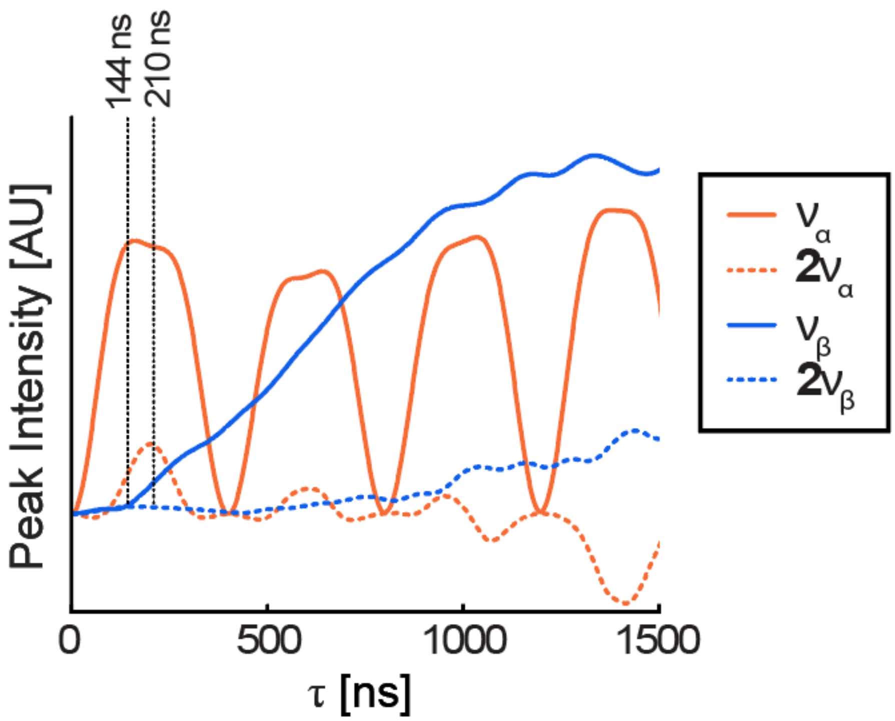

**Figure S5. Simulation of [15]N ESEEM frequency peaks intensities with respect to τ.** Relative peak intensities for the $\nu_\alpha$, $\nu_\beta$, $2\nu_\alpha$, and $2\nu_\beta$ transitions calculated from Equation S11 at varying τ. The vertical lines represent the τ explored experimentally. Here, τ = 144 ns gives a high intensity $\nu_\alpha$ but $\nu_\beta$ is negligible. This agrees with experimental results outlined in Figure S6. At τ = 210 ns, we anticipate a similar intensity of $\nu_\alpha$, with a small $\nu_\beta$ and $2\nu_\alpha$ peak. $2\nu_\beta$ cannot be observed until much longer τ. However, $T_m$ relaxation would significantly influence the intensity of the ESEEM echo at these values.

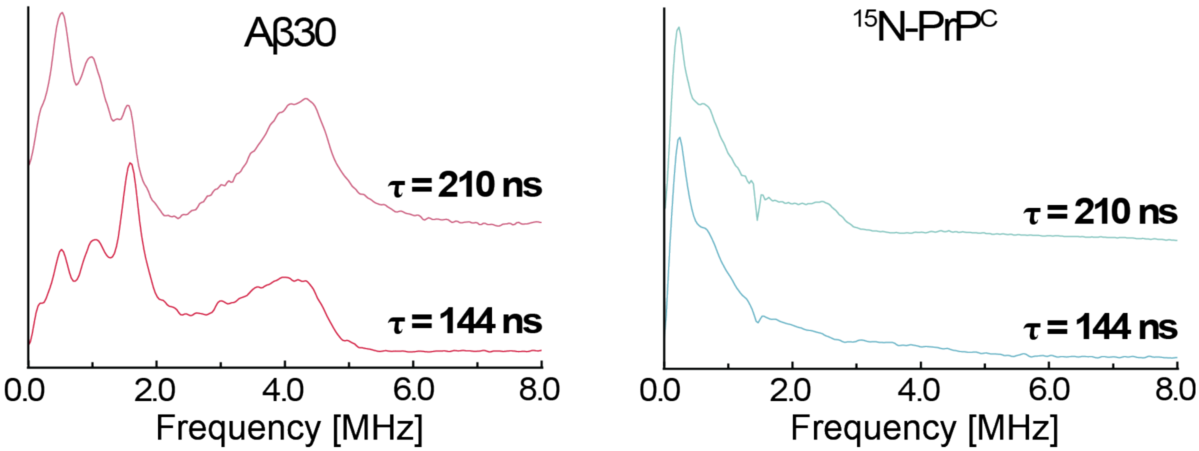

**Figure S6. Comparison between τ values in ESEEM frequency domain of $^{14}N$ and $^{15}N$.** ESEEM frequency domain of Aβ30 (left) and $^{15}N$-$PrP^C$ (right) show the comparison of samples obtained at a τ of 210 ns (top) and 144 ns (bottom), T initially equaling 12 ns. These τ were chosen to suppress the $^1H$ signal at 3316 G. τ = 144 ns slightly distorts the $^{14}N$ peaks and does not resolve the $\nu_\beta$ for $^{15}N$, as predicted by Figure S5. Thus, τ = 210 ns was used for subsequent experiments.

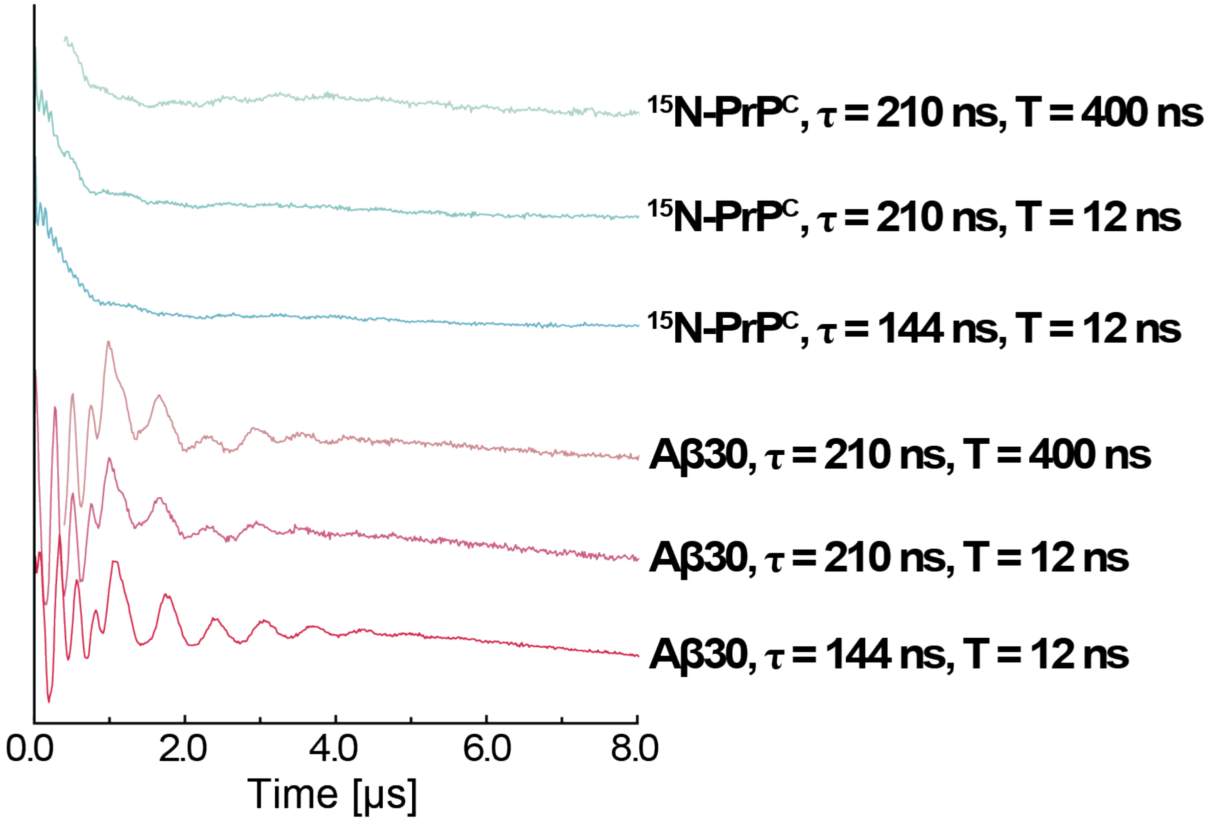

**Figure S7. Experimental ESEEM raw time domain signal.** ESEEM time domain signal of $^{15}$N-PrP$^{C}$ (blue) and Aβ30 (red) at different τ and initial T values. T and τ values are reported with their respective time domain signals. The shortest T was 12 ns as that is the lower limit without significant distortion due to pulse overlap.

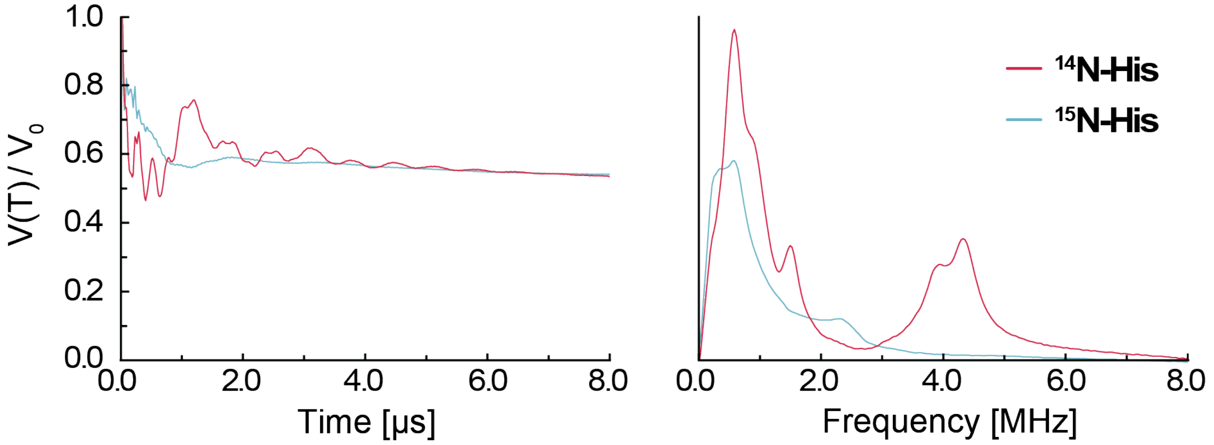

**Figure S8. ESEEM signal intensity comparison of $^{14}N$ and $^{15}N$.** ESEEM time domain (left) and frequency domain (right) of $^{14}N$ and $^{15}N$ histidine in complex with $Cu^{2+}$. The samples were prepared with a 1:4 ratio of $Cu^{2+}$ : $^{14}N$ or $^{15}N$ histidine, with a $Cu^{2+}$ concentration of 300 μM. The time domain signals were normalized to V(0) = 1 to account for subtle variations in sample prep, spectrometer tuning, and spectral changes between the two systems. The $^{14}N$ nuclei yield a greater peak intensity compared to the $^{15}N$ nuclei

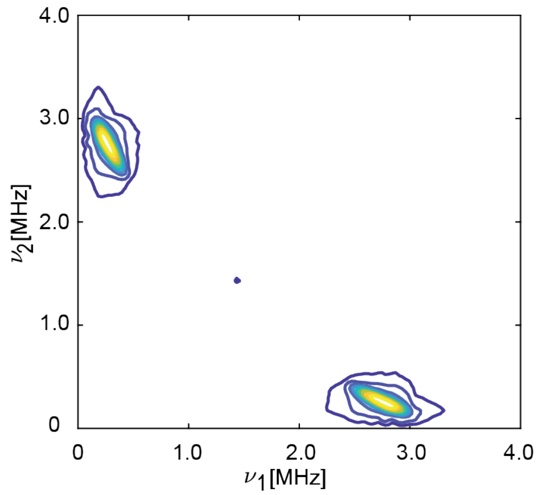

**Figure S9. Intensity comparison between strongly and weakly coupled $^{15}$N.** Simulated HYSCORE spectrum of $Cu^{2+}$ interacting with one coupled and five weakly coupled $^{15}$N. Despite there being 5x weakly coupled $^{15}$N, the coupled $^{15}$N signal intensity if much larger. The strongly coupled $^{15}$N has an $A_{xx} = A_{yy} = 3.2$ MHz, $A_{zz} = 2.0$ MHz, $A_{iso} = 2.8$ MHz, T = -0.4 MHz and weakly coupled $A_{xx} = A_{yy} = 0.1$ MHz, $A_{zz} = 0.01$ MHz, $A_{iso} = 0.07$ MHz, T = -0.03 MHz. The small amount of anisotropy added to the weakly coupled hyperfine tensor was implemented to maximize its peak intensity without causing any observable splitting. Notably, the experimentally observed peak is significant. This is likely a consequence of one of two possibilities.

1) The peak might be due to coupling with a specific $^{15}$N within $PrP^C$, giving rise to some nonnegligible, yet unquantifiable anisotropy, thus producing a significant signal.
2) There is an interaction with the bath of $^{15}$N, therefore stemming from the contribution from many weakly coupled $^{15}$N nuclei.

Since the number of nuclei does not influence peak position or shape for weakly coupled $^{15}$N, as shown in Figure S10, both are logical conclusions.

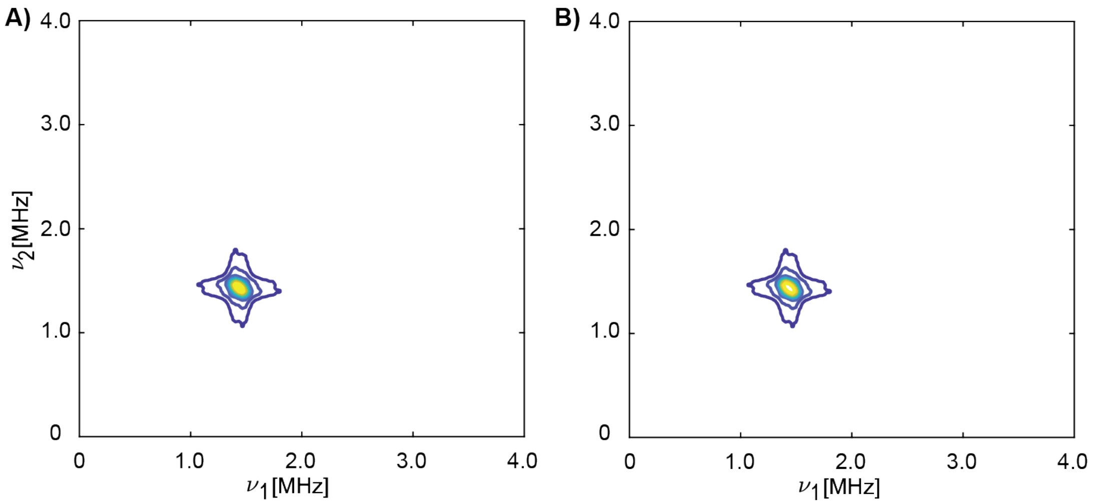

**Figure S10. Effect of multiple weakly coupled $^{15}$N nuclei in HYSCORE.** Simulated HYSCORE spectra of A) one and B) six weakly coupled $^{15}$N. The number of weakly coupled $^{15}$N has no observable effect on the simulated spectra. $A_{xx} = A_{yy} = 0.1$ MHz, $A_{zz} = 0.01$ MHz, $A_{iso} = 0.07$ MHz, T = -0.03 MHz. The small amount of anisotropy added to the weakly coupled hyperfine tensor was implemented to maximize the intensity of the weakly coupled peak without causing any observable splitting. Here, we observe a single peak along the diagonal at (1.45 MHz, 1.45 MHz). In comparison to the experimental HYSCORE, this peak lines up with the (1.47 MHz, 1.47 MHz) peak, confirming this peak is due to weakly coupled nuclei. Additionally, the presence of multiple weakly coupled nuclei does not influence peak position or shape. This is likely due to the low dipolar coupling, T, as shown in Figure S12.

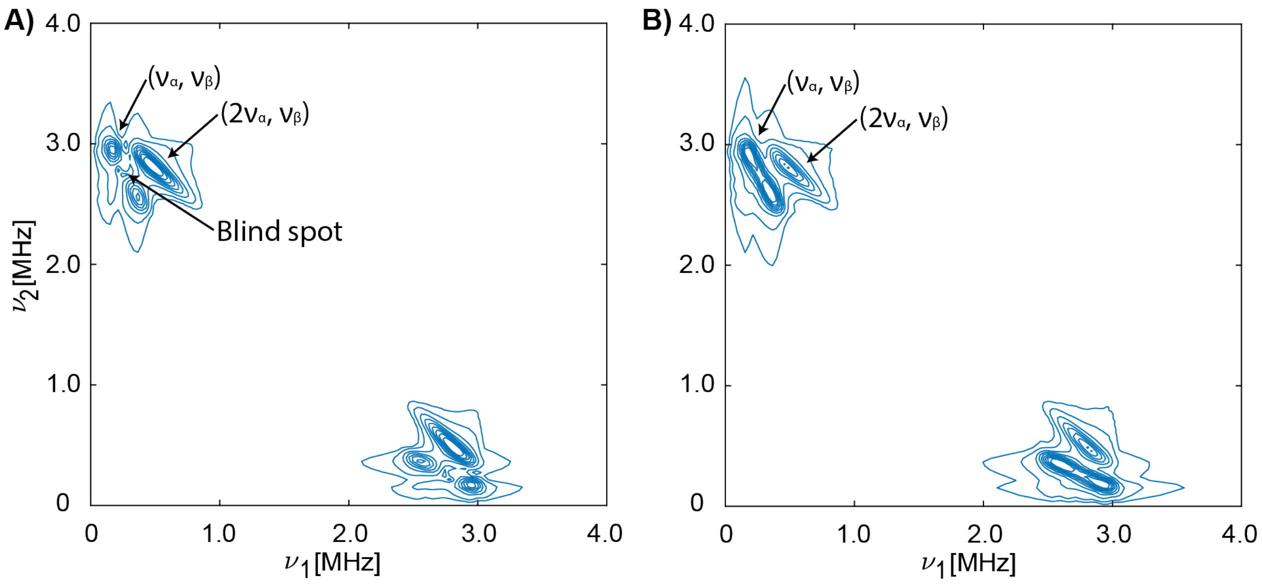

**Figure S11. Blindspot in HYSCORE simulations.** Simulated HYSCORE spectrum A) with $\tau$ = 210 ns and B) implementing $\tau$ averaging from $\tau$ = 100 to 400 ns in 10 ns steps. Without $\tau$ averaging, blind spots are possible. However, the frequencies of either of the two ($2\nu_\alpha$, $\nu_\beta$) peaks resulting from the blind spot split do not match that observed for the ($2\nu_\alpha$, $\nu_\beta$) peak. With two $^{15}N$ and utilizing $\tau$ averaging, the resulting HYSCORE simulation best represents the experimental $^{15}N$-$PrP^C$ in Figure 3.

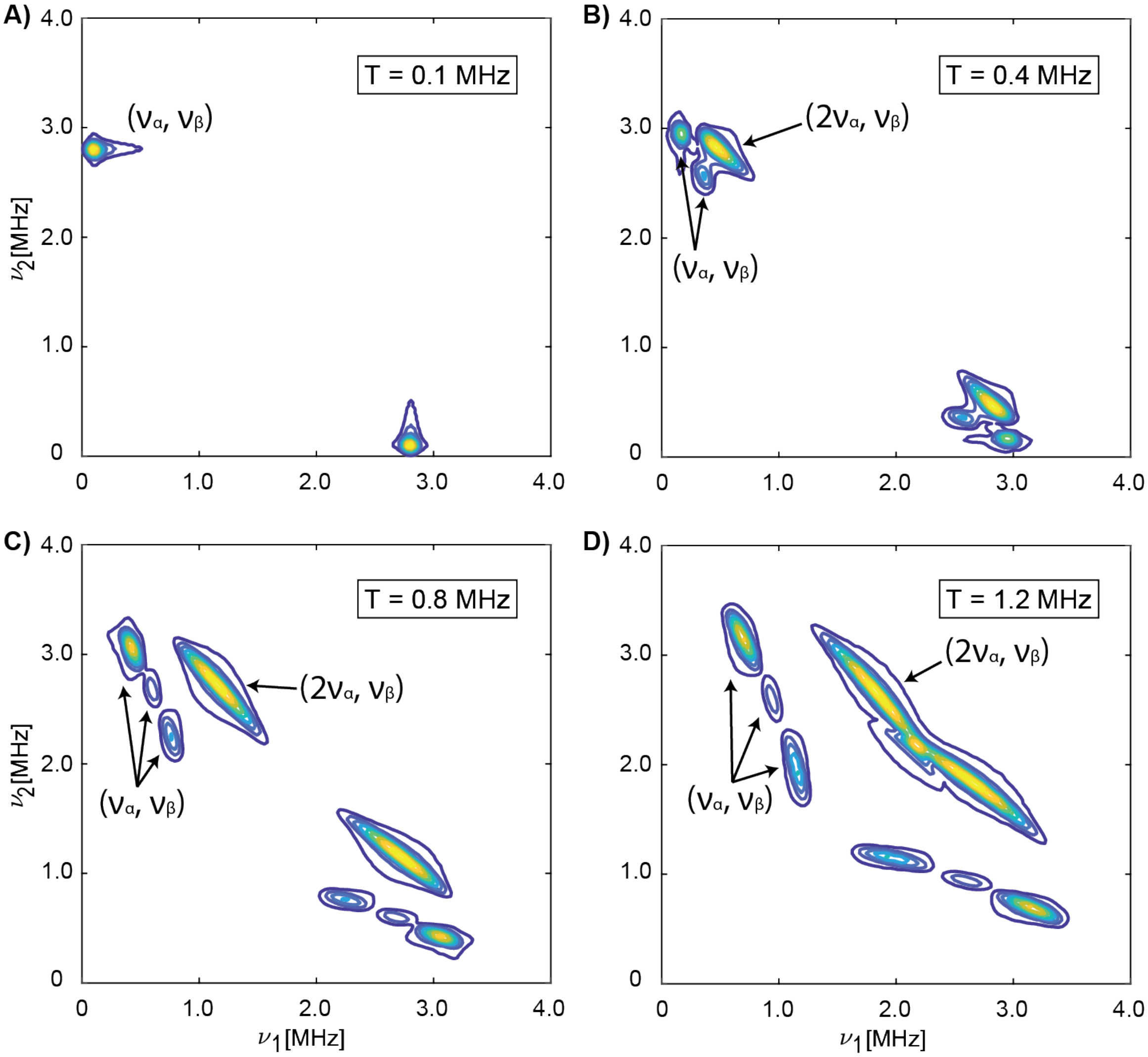

**Figure S12. HYSCORE simulation with varying dipolar coupling.** Simulated HYSCORE of two $^{15}N$ nuclei with $A_{iso}$ = 2.8 MHz, τ = 210 ns, A) T = -0.1 MHz, B) T = -0.4 MHz, C) T = -0.8 MHz, and D) T = -1.2 MHz. The gaps in the ($\nu_\alpha$, $\nu_\beta$) cross peak are due to τ blind spots. For T = -0.1 MHz, only the ($\nu_\alpha$, $\nu_\beta$) cross peak appears.

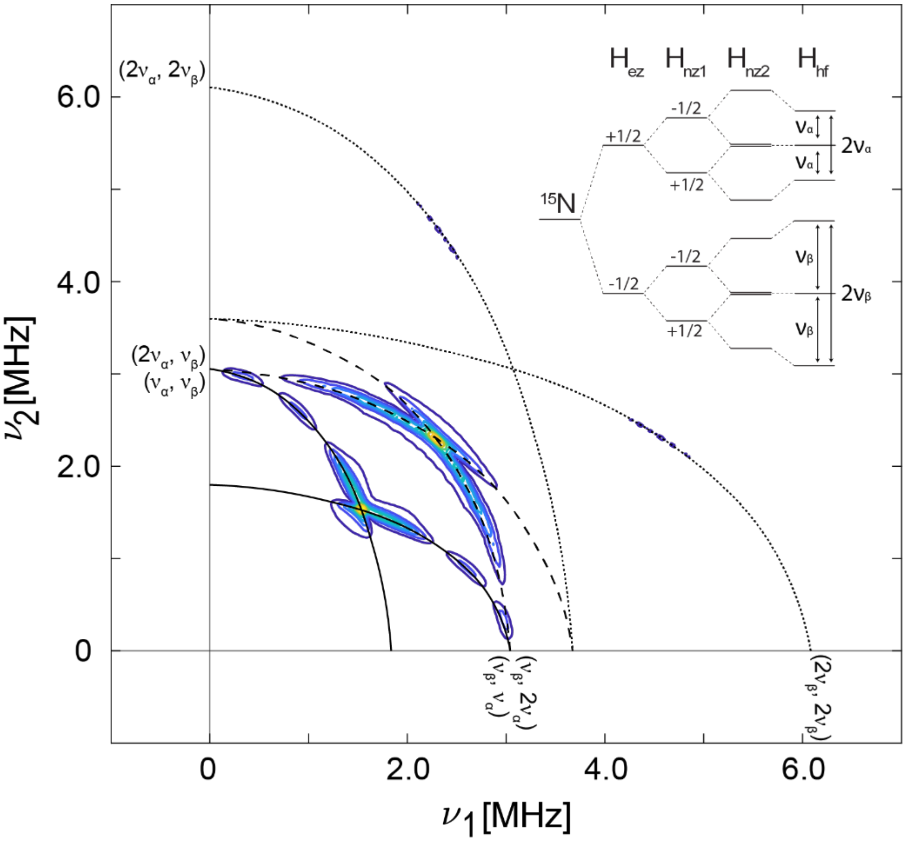

**Figure S13. HYSCORE simulation with large dipolar coupling.** Simulated HYSCORE of two $^{15}N$ nuclei with $A_{iso}$ = 2.0 MHz, T = -1.2 MHz, $\tau$ = 210 ns. The black solid line was set to overlay on the ($\nu_\alpha$, $\nu_\beta$) cross peak. The dashed line is set to exactly twice $\nu_1$ of the solid line, i.e., would expect to overlay with the ($2\nu_\alpha$, $\nu_\beta$) cross peak. Similarly, the dotted line is at twice $\nu_1$ and twice $\nu_2$ of the solid line, thus should overlay with ($2\nu_\alpha$, $2\nu_\beta$). The diagonally symmetric peaks are also fit in a similar manner. The inset contains the energy level diagram for a two $^{15}N$ system. Vertical lines indicate the transitions observed in this HYSCORE simulation. By incorporating a much larger dipolar coupling in this simulation, in comparison to the T = -0.4 MHz used in Figure 3 and S11, the ($2\nu_\alpha$, $\nu_\beta$) cross peak is more prominent and a new peak appears at ($2\nu_\alpha$, $2\nu_\beta$). Thus, it is likely these new peaks are dependent both on the presence of two $^{15}N$ nuclei and a large dipolar coupling. The multiple peaks within the ($\nu_\alpha$, $\nu_\beta$) cross peak are due to blind spots.

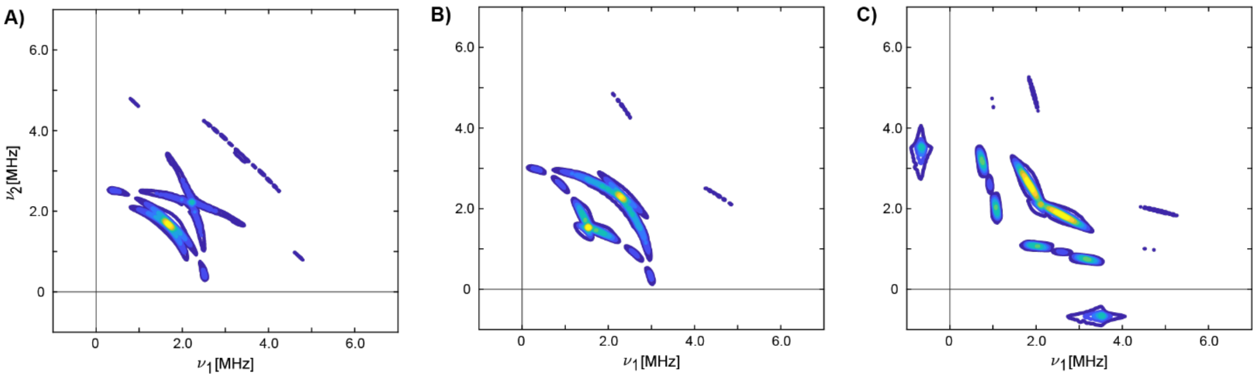

**Figure S14. HYSCORE simulation with varying hyperfine coupling.** Simulated HYSCORE spectra with an isotropic hyperfine coupling, $A_{iso}$, of A) 1.0 MHz, B) 2.0 MHz, and C) 3.0 MHz. The dipolar coupling is held constant at T = -1.2 MHz. From this, we see that at high dipolar coupling, the cross peaks at ($2\nu_\alpha$, $\nu_\beta$) and ($2\nu_\alpha$, $2\nu_\beta$) remain regardless of the $A_{iso}$ used. Thus, these transitions are likely accessible primarily due to larger anisotropy, i.e., larger T, rather than a large $A_{iso}$.

**Expressions for calculating ESEEM spectra.**

Equations S1 and S2 represent the time domain signal for a primary echo, V(τ) from 2-pulse ESEEM, for a single I = ½ nucleus. For two I = ½ nuclei, the resulting signal follows Equation S3, which can be simplified to the square of equation 7 when the nuclei are equivalent.[2]

$$V_{I=\frac{1}{2}}(\tau) = 1 - \frac{k}{2}A(\tau) \quad \textbf{(S1)}$$

$$A(\tau) = 1 - \cos\omega_\alpha\tau - \cos\omega_\beta\tau + \frac{1}{2}\cos(\omega_\alpha + \omega_\beta)\tau + \frac{1}{2}\cos(\omega_\alpha - \omega_\beta)\tau \quad \textbf{(S2)}$$

$$V_{I_1=\frac{1}{2},I_2=\frac{1}{2}}(\tau) = V^2_{I=\frac{1}{2}}(\tau) = 1 - kA(\tau) + \frac{k^2}{4}A^2(\tau) \quad \textbf{(S3)}$$

$$k = \left(\frac{B\omega_{nz}}{\omega_\alpha\omega_\beta}\right)^2 \quad \textbf{(S4)}$$

$$B = 3Tcos\theta sin\theta \quad \textbf{(S5)}$$

Here, k is the modulation depth parameter dependent on the dipolar coupling, ω is equal to 2πν, τ is the pulse timing, T is the dipolar coupling, and θ is the angle between the electron-nuclear vector and the direction of the applied magnetic field, $B_0$. Upon a trigonometric expansion of Equation S3, a set of double frequency terms arise as shown in Equation S6. A full expansion is shown in Equation S7.

$$V_{DF}(\tau) = \frac{3k^2}{16}\left[1 + \cos 2\omega_\alpha\tau + \cos 2\omega_\beta\tau + \frac{1}{6}\cos(2\omega_\alpha + 2\omega_\beta)\tau + \frac{1}{6}\cos(2\omega_\alpha - 2\omega_\beta)\tau\right] \quad \textbf{(S6)}$$

It is evident from the modulation depth, $3k^2/16$, that the intensity of the double frequency transition is highly dependent on the dipolar coupling.

$$V_{I_1=\frac{1}{2},I_2=\frac{1}{2}}(\tau) = 1 - k\left[1 - \cos\omega_\alpha\tau - \cos\omega_\beta\tau + \frac{1}{2}\cos(\omega_\alpha + \omega_\beta)\tau + \frac{1}{2}\cos(\omega_\alpha - \omega_\beta)\tau\right] \quad \textbf{(S7)}$$

$$+ \frac{k^2}{4}\left[1 - 3\cos\omega_\alpha\tau - 3\cos\omega_\beta\tau + 2\cos(\omega_\alpha + \omega_\beta)\tau + 2\cos(\omega_\alpha - \omega_\beta)\tau\right]$$

$$+ \frac{3k^2}{16}\left[1 + \cos 2\omega_\alpha\tau + \cos 2\omega_\beta\tau + \frac{1}{6}\cos(2\omega_\alpha + 2\omega_\beta)\tau + \frac{1}{6}\cos(2\omega_\alpha - 2\omega_\beta)\tau\right]$$

$$+ \frac{k^2}{8}\left[1 - \cos(2\omega_\alpha + \omega_\beta)\tau - \cos(2\omega_\alpha - \omega_\beta)\tau - \cos(\omega_\alpha + 2\omega_\beta)\tau - \cos(\omega_\alpha - 2\omega_\beta)\tau\right]$$

Equations S8 and S9 represent the time domain signal for a stimulated echo, V(τ,T) from a 3-pulse ESEEM, for a single I = ½ nucleus. For two, equivalent I = ½ nuclei, the resulting signal follows Equation S10.

$$V_{I=\frac{1}{2}}(\tau, T) = \frac{1}{2}\left[V_\alpha(\tau, T) + V_\beta(\tau, T)\right] \quad \textbf{(S8)}$$

$$V_\alpha(\tau, T) = 1 - \frac{k}{2}\left[(1 - \cos\omega_\beta\tau)(1 - \cos\omega_\alpha(\tau + T))\right] \quad \textbf{(S9)}$$

$$V_\beta(\tau, T) = 1 - \frac{k}{2}\left[(1 - \cos\omega_\alpha\tau)(1 - \cos\omega_\beta(\tau + T))\right] \quad \textbf{(S9')}$$

$$V_{I_1=\frac{1}{2},I_2=\frac{1}{2}}(\tau, T) = \frac{1}{2}\left[V_\alpha^2(\tau, T) + V_\beta^2(\tau, T)\right] \quad \textbf{(S10)}$$

Upon a trigonometric expansion of Equation S10, we obtain S11.

$$V_{I_1=\frac{1}{2},I_2=\frac{1}{2}}(\tau,T) = \frac{1}{2}\left[1 - k(1-\cos\omega_b\tau) + \frac{k^2}{4}\left(\frac{9}{4} - 3\cos\omega_b\tau + \frac{3}{4}\cos 2\omega_b\tau\right)\right. \quad \textbf{(S11)}$$

$$+k\left((1-\cos\omega_b\tau) - \frac{k}{4}(3 - 4\cos\omega_b\tau + \cos 2\omega_b\tau)\right)\cos\omega_\alpha(\tau+T)$$

$$\left.+\frac{k^2}{16}\left(3 - 4\cos\omega_b\tau + \cos 2\omega_\beta\tau\right)\cos 2\omega_\alpha(\tau+T)\right]$$

$$+\frac{1}{2}\left[1 - k(1-\cos\omega_\alpha\tau) + \frac{k^2}{4}\left(\frac{9}{4} - 3\cos\omega_\alpha\tau + \frac{3}{4}\cos 2\omega_\alpha\tau\right)\right.$$

$$+k\left((1-\cos\omega_\alpha\tau) - \frac{k}{4}(3 - 4\cos\omega_\alpha\tau + \cos 2\omega_\alpha\tau)\right)\cos\omega_\beta(\tau+T)$$

$$\left.+\frac{k^2}{16}(3 - 4\cos\omega_\alpha\tau + \cos 2\omega_\alpha\tau)\cos 2\omega_\beta(\tau+T)\right]$$
